# MASTR-seq: Multiplexed Analysis of Short Tandem Repeats with sequencing

**DOI:** 10.1101/2024.04.29.591790

**Authors:** Chuanbin Su, Keerthivasan Raanin Chandradoss, Thomas Malachowski, Ravi Boya, Han-Seul Ryu, Kristen J. Brennand, Jennifer E. Phillips-Cremins

**Affiliations:** Department of Bioengineering, University of Pennsylvania, Philadelphia, PA; Epigenetics Institute, Perelman School of Medicine, University of Pennsylvania, Philadelphia, PA; Department of Genetics, Perelman School of Medicine, University of Pennsylvania, Philadelphia, PA; Nash Family Department of Neuroscience, Icahn School of Medicine at Mount Sinai, New York, NY 10029; Department of Genetics and Genomics, Icahn School of Medicine at Mount Sinai, New York, NY 10029; Friedman Brain Institute, Black Family Stem Cell Institute, Pamela Sklar Division of Psychiatric Genomics, Icahn School of Medicine at Mount Sinai, New York, NY 10029; Department of Psychiatry, Yale School of Medicine, New Haven, CT, 06520

## Abstract

More than 60 human disorders have been linked to unstable expansion of short tandem repeat (STR) tracts. STR length and the extent of DNA methylation is linked to disease pathology and can be mosaic in a cell type-specific manner in several repeat expansion disorders. Mosaic phenomenon have been difficult to study to date due to technical bias intrinsic to repeat sequences and the need for multi-modal measurements at single-allele resolution. Nanopore long-read sequencing accurately measures STR length and DNA methylation in the same single molecule but is cost prohibitive for studies assessing a target locus across multiple experimental conditions or patient samples. Here, we describe MASTR-seq, <u>M</u>ultiplexed <u>A</u>nalysis of <u>S</u>hort <u>T</u>andem <u>R</u>epeats, for cost-effective, high-throughput, accurate, multi-modal measurements of DNA methylation and STR genotype at single-allele resolution. MASTR-seq couples long-read sequencing, Cas9-mediated target enrichment, and PCR-free multiplexed barcoding to achieve a >ten-fold increase in on-target read mapping for 8-12 pooled samples in a single MinION flow cell. We provide a detailed experimental protocol and computational tools and present evidence that MASTR-seq quantifies tract length and DNA methylation status for CGG and CAG STR loci in normal-length and mutation-length human cell lines. The MASTR-seq protocol takes approximately eight days for experiments and one additional day for data processing and analyses.

**Key points:**

- We provide a protocol for MASTR-seq: <u>M</u>ultiplexed <u>A</u>nalysis of <u>S</u>hort <u>T</u>andem <u>R</u>epeats using Cas9-mediated target enrichment and PCR-free, multiplexed nanopore sequencing.
- MASTR-seq achieves a >10-fold increase in on-target read proportion for highly repetitive, technically inaccessible regions of the genome relevant for human health and disease.
- MASTR-seq allows for high-throughput, efficient, accurate, and cost-effective measurement of STR length and DNA methylation in the same single allele for up to 8-12 samples in parallel in one Nanopore MinION flow cell.

## Introduction

More than sixty human diseases have been linked to the mutation-length expansion of short tandem repeat (STR) tracts^1-6^. Accurate genotyping of intractable repetitive sequences has thus far been limited by several technical challenges^7,8^. First, tandem repeat DNA sequences slip during PCR amplification resulting in technical biases in quantifying amplicon length. Second, short-read sequencing fails to span mutation-length STRs in many cases, thus prohibiting direct measurement of the DNA sequence and underestimating the degree of mosaicism of STR expansion in disease. Third, challenges arise in computational methods to map short reads over repeat regions, and they’re generally removed for downstream analysis^7, 9, 10^. Thus, there is a need for technology advances that allow for accurate and cost-effective approaches for sequencing STR tracts in patient-derived samples with unknown genotypes.

Long-read sequencing provides a significant technical advance to overcome the challenges prohibiting direct genotyping of large copy number variations in the repetitive genome. Whole genome long-read sequencing was used for the accurate assembly of telomere-to-telomere reference genome maps from haploid and diploid cell lines^11-14^. It has also been effectively applied to accurately genotype a range of repetitive sequences, including STR tracts, short interspersed nuclear elements (SINEs), long interspersed nuclear elements (LINEs), centromeres, pericentromeric regions, and telomeres^15-26^. However, for studies focused on genotyping a specific locus across a number of samples and conditions, genome-wide long-read sequencing suffers from low sample throughput, low on-target coverage, and the prohibitively high financial burden to achieve the locus-specific read coverage required to accurately measure STR mosaics^18, 27-31^. Approaches to capture a predetermined target locus and sequence with nanopore long-reads have been effectively used to genotype structural variants and STR tracts^18, 28, 32-36^. However, targeted long-read sequencing methods remain limited by low sample throughput, high off-target nonspecific background reads, low on-target locus-specific read coverage, and prohibitive cost.

Recently, multiple strategies for creating efficient targeted long-read methods have been proposed. Attempts to reduce background reads in hybrid capture or Cas9-mediated targeting approaches might involve using exonuclease digestion or affinity-based removal of Cas9 bound non-target DNA^37-42^. Real-time target enrichment methods to circumvent a reagent-based molecular targeting step have also been developed, including Read Until^43-45^, UNCALLED^46^ and ReadFish^47^. Undesired DNA molecules are removed from the pore by reversing the current in the channel, thus allowing only targeted DNA molecules to be fully sequenced without additional specialized reagents^43-47^. Nevertheless, limitations in multiplexed sample throughput and low signal-to-noise read specificity remain, thus emphasizing the need for a cost-effective, high-throughput, on-target long-read enrichment method to accurately measure defined STR tracts across multiple samples.

To address these limitations, we developed MASTR-seq - <u>M</u>ultiplexed <u>A</u>nalysis of <u>S</u>hort <u>T</u>andem <u>R</u>epeats - as a Cas9-mediated, size selection-based, PCR-free long-read sequencing method for multi-modal, multiplexed measurement of genotype and DNA methylation at a predetermined STR tract. MASTR-seq aims to minimize genomic background while preserving the integrity of on-target genomic fragments. Upon Cas9-mediated target enrichment with multiple guide RNAs^33^, we add a fragment size selection step and multiplexed PCR-free barcoding to increase the proportion of on-target reads spanning the full mutation-length STR, thus enabling the pooling and multi-modal long-read sequencing of 8-12 independent samples in parallel.

### Overview of MASTR-seq

MASTR-seq is a multiplexed, increased sample throughput, on-target nanopore long-read sequencing method designed for multi-modal measurement of repeat genotype and DNA methylation in the same single allele. MASTR-seq is particularly useful for loci containing intractable mutation-length STRs in complex mosaic patient-derived cell lines and samples. The protocol involves five major stages (**Figure 1**): (Day 1-2) High molecular weight (HMW) genomic DNA (gDNA) preparation, (Day 3) Cas9-mediated target DNA cleavage and dA tailing, (Day 4-5) Target DNA fragment size selection, (Day 6) PCR-free library preparation via multiplexed barcoding, and (Day 7-9) Sequencing and data analysis.

**Figure 1.**
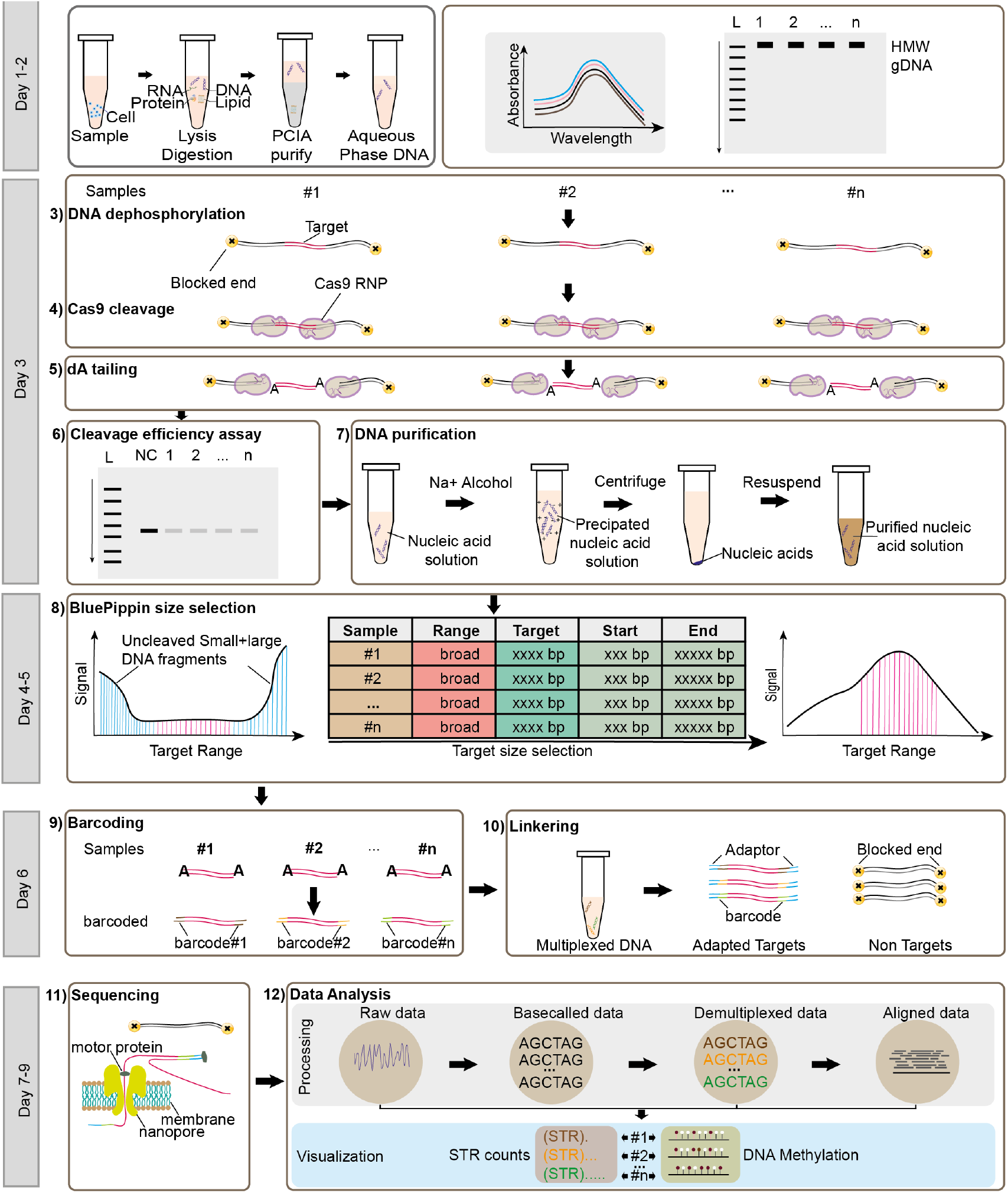
Schematic of MASTR-seq procedure. **Days 1-2:** (step 1) Samples are trypsinized to single-cell suspension and cells are lysed. RNA and protein are digested by RNase A and Proteinase K, respectively. DNA is subsequently purified by phenol:chloroform:isoamyl alcohol (PCIA, 25:24:1). (step 2) HMW genomic DNA quality control assays are performed using Nanodrop for DNA quality, Qubit for quantifying DNA concentration, and gel electrophoresis for DNA integrity. **Day 3**: crRNA/tracrRNA annealing and Cas9 RNP assembling are completed (not shown in this schematic) before (step 3) HMW DNA is dephosphorylated, (step 4) Cas9-mediated RNP cleavage, and (step 5) dA-tailing. (step 6) The quality of Cas9-mediated RNP cleavage efficiency is assessed by PCR amplification of the target region using flanking primers. (step 7) Cas9 RNP cleaved products are purified from salts using sodium acetate-ethanol precipitation. **Days 4-5:** (step 8) Target DNA fragments are precisely size selected using BluePippin. **Day 6:** (step 9) Samples are barcoded in a PCR-free manner and pooled for multiplexing using nanopore native barcodes. (step 10) Adapters are ligated onto the multiplexed DNA. **Days 7-9:** (step 11) DNA is loaded onto the Nanopore flow cell and sequenced for 48-72 hours. (step 12) Raw data is processed by basecalling, demultiplexing, and aligning sequentially. Custom computational tools are employed for the analyses of single-allele DNA methylation and STR tract length.

### (Day 1-2) High molecular weight (HMW) genomic DNA (gDNA) preparation

1. Sample preparation: Cells are harvested using line-specific single cell dissociation reagents.
2. Cell counting normalization: Cell numbers are determined using an automated Countess system or manual counting with a hemocytometer.
3. HMW gDNA extraction: HMW gDNA is extracted using phenol:chloroform:isoamyl alcohol (PCIA, 25:24:1) extraction followed by ethanol precipitation.
4. HMW gDNA quality control (QC) assay: DNA quality and concentration are assessed using Nanodrop and Qubit and DNA integrity is verified using a 0.5% agarose gel.

### (Day 3) Cas9-mediated target DNA cleavage and dA tailing

1. CRISPR RNA (crRNA) annealing: Equimolar crRNAs and trans-activating crRNA (tracrRNA) are pooled and annealed in a 1:1 molar ratio.
2. Cas9 and crRNA complex formation: Annealed crRNAs and tracrRNA (1:1) are assembled with Cas9 to form Cas9 ribonucleoprotein (RNP) complexes.
3. HMW genomic DNA dephosphorylation: NEB Quick calf intestinal alkaline phosphatase (CIP) is used to dephosphorylate DNA.
4. Cas9 cleavage and dA-tailing: Cas9 RNP complex cleaves dephosphorylated DNA, followed by Taq polymerase-mediated dA tailing to the 3’ end of dephosphorylated DNA.
5. Cas9-mediated cleavage quality control check: A PCR reaction is performed to confirm Cas9 RNP cleavage efficiency using target flanking primers.
6. DNA purification: Cas9 RNP digested products are precipitated using ethanol. Precipitated DNA is washed by 75% ethanol and rehydrated in 10 mM TRIS-HCL (PH 8.0).

### (Day 4-5) Target DNA fragment size selection

1. BluePippin is utilized for size selection to remove uncleaved or non-target size DNA fragments.

### (Day 6) PCR-free library preparation via multiplexed barcoding

1. Multiplexed barcoding: Samples are multiplexed via PCR-free ligation of nanopore barcodes.
2. Adapter ligation: Adapters are ligated to the barcoded multiplexed samples.

### (Day 7-9) Sequencing and data analysis

1. Loading library: Library DNA is mixed with sequencing buffers and loading beads.
2. Sequencing: Nanopore sequencing is set up on the lab installed MinKnow platform.
3. Basecalling: Guppy (V6.2.1) is used for base calling to convert raw electrical signals to quality sequencing reads in .fastq format.
4. Debarcoding: Guppy (V6.2.1) is used for demultiplexing reads into unique sample.
5. Alignment: Guppy (V6.2.1) is used for mapping demultiplexed reads to the reference genome.
6. Visualization: Customized computational tools are employed for the visualization of STR genotype and DNA methylation.

### Advantages & Applications

In our recently published work focused on fragile X syndrome (FXS)^48, 49^, we applied our method to accurately measure CGG STR length and DNA methylation for the 5’UTR and promoter of the *FMR1* gene in the same single-allele across several pooled normal-length, premutation-length, and mutation-length patient-derived iPSC lines. By enriching the proportion of long-reads spanning the target locus by >10-fold compared to the nanopore Cas9-targeted sequencing (nCAT) method^33^, we achieved >300x on-target read coverage at the *FMR1* CGG STR in a pool of 8-12 independent samples in a single MinION flow cell (BOX6). We also demonstrate here that MASTR-seq is effective at measuring the CAG STR in the *HTT* gene in human embryonic stem cell lines after knock-in of specific mutation-length CAG tracts. The ability to precisely measure the number of repeat units at single-allele resolution represents a major step change improvement in sensitivity and specificity from traditional gold-standard STR genotyping methods used in the clinic such as triplet repeat primed PCR (TP-PCR) (AmplideX, Asuragen). Moreover, MASTR-seq offers repeat unit resolution and single-molecule measurements that cannot be achieved by traditional Southern blots. The MASTR-seq protocol can be easily adapted for a wide range of applications for multiplexed, cost-effective, multi-modal analysis of repeat genotype and DNA methylation at single-allele resolution. MASTR-seq future applications include:

### Multi-modal analysis of intractable repetitive sequences for genotype and DNA methylation

Long-read sequencing has been successfully applied for the detection of copy number variants and DNA methylation of repetitive sequences, including STRs, transposable elements^18, 20, 28, 32, 34, 48^, centromeres^50^, peri-centromeres, and telomeres^51, 52^ across a wide range of samples. Nanopore long-reads can simultaneously detect mosaic DNA methylation and somatic sequence variation^53,54^. Here, we demonstrate that MASTR-seq provides a facile cost-effective way to increase on-target reads to quantify repeat unit length and DNA methylation in the same single-read in pre-determined repetitive sequences across multiple samples at a single-allele resolution. MASTR-seq will broadly scale to applications in which there is a specific repetitive sequence of interest across multiple samples to understand genotype and DNA methylation variation. The applications are not limited to STR tracts, and could involve genotyping technically intractable SINE, LINE, centromere, and telomere tracts for a high-throughput comparison of multiple samples in one nanopore run.

### Multiplexed PCR-free barcoding for high-throughput pooling and cost-effective multi-modal measurements as a defined target locus in multiple samples

Previously published methods such as nanopore Cas9-targeted sequencing (nCAT)^33^ highlighted the power of the Cas9-mediated enrichment step in allowing the pooling of up to 10 target loci from the same single cell line in one flow cell. Here, by adding a large fragment size selection step, we increase on-target read proportion >10-fold for a single locus compared to published nCAT data (BOX6). We further employ PCR-free multiplexed barcoding to allow for the pooling of up to 8-12 different patient-derived cell line samples in a single MinION flow cell.

### Comparison with other methods

Over the last few years, several variations on long-read sequencing have been developed to enrich for specific genomic DNA sequences^18, 28, 32-35^. MASTR-seq builds on the Cas9-mediated enrichment approach by adding several features useful for basic science and clinical applications. nCAT shows an ∼0.5% on-target read mapping rate for multiple pooled targets from a single sample even despite the Cas9-mediated enrichment step, thus highlighting the need to further improve the cost, specificity, and throughput of the method.

We add a long fragment size selection step to increase the on-target read proportion by >10-fold, therefore allowing for the re-distribution of the read availability to allow for multiplexing multiple samples. We also add PCR-free multiplexed barcoding to pool multiple samples and achieve >300x on-target read coverage for the target locus for 8-12 samples from a single MinION flow cell. The presence of off-target DNA fragments results in competition of reads during nanopore sequencing, leading to low on-target read yield. Prior to the improvements of MASTR-seq, only a very limited number of samples could be sequenced in parallel due to severe challenges in cost and poor sensitivity in genotype measurements due to low on-target read coverage. MASTR-seq’s advances enable significant decreases in cost for multi-modal measurements of target regions across multiple samples. When quantifying STR tract length, it is preferable to use a method with lower susceptibility to amplification biases. MASTR-seq is a Cas9-mediated nanopore long-read sequencing approach that does not involve PCR amplification steps during library preparation. A PCR-free sequencing approach offers the advantage of reducing errors by eliminating polymerase-based errors that occur at high frequency in repetitive regions of the genome. Finally, we present a simplified yet optimized computational pipeline which enables researchers to accurately analyze STR length and DNA methylation in the same single molecules with minimal new technology development.

### Limitations and possible solutions

MASTR-seq requires over 5 μg DNA input to achieve efficient on-target coverage. When using rare tissue samples, there is limited DNA amount available. Potential solutions to improve library preparation efficiency for limited DNA input, include: 1) streamlining the workflow with library adapter linked barcodes for one-step ligation; 2) exploring alternative methods of target enrichment, such as oligo hybridization-based capture instead of Cas9-mediated enrichment; 3) increasing the efficiency of the DNA extraction, Cas9 cleavage, and DNA size selection steps, or 4) merging DNA extraction, Cas9 cleavage and size selection together as mentioned in the Cas9-Assisted Targeting of CHromosome segments(CATCH) method^55^. The BluePippin size selection step currently takes at least 6 hours, depending on sample quantity. To address this issue, it might be possible in the future to use the PippinHT System’s high-throughput solution, which streamlines the process and achieves sample reproducibility within 6 hours. This system has a capacity for 24 samples, significantly reducing time investment and meeting the demands of high-throughput-sequencing. Moreover, in our current MASTR-seq protocol, we use the MinION flow cell (R9.4.1) paired with Guppy high-accuracy basecalling to generate sequences with an accuracy of 97.6% and a Phred score of Q16^56-58^. However, our methods are still limited in long-read accuracy, which might be improved in the future by implementing an upgraded Q20+ enzyme of V14 chemistry, in conjunction with the new flow cell (R10.4.1), or utilizing the latest base-calling algorithm dorado (https://github.com/nanoporetech/dorado).

### Experimental design

#### CRISPR RNA (crRNA) design

We used the CHOPCHOP tool to design distinct crRNAs targeting sequences flanking the target locus (**BOX 1**). The parameters used were: In: GRCh38, Target: *FMR1* or *HTT* genomic coordinates, For: nanopore enrichment, Using: CRISPR/Cas9. To increase on-target coverage, we designed at least 2 crRNAs upstream and downstream of the targets as reported previously^33^. We refined the selection of candidate crRNAs based on several parameters: cleavage location, (+) strand crRNAs for the region upstream of the locus, (-) strand crRNAs for the region downstream of the locus, 40-80% GC content, self-complementary=0, mismatches (MM0=0; MM1=0; MM2<2), and cleavage efficiency>50. Final crRNA DNA sequences were synthesized by Integrated DNA Technologies and the sequences are provided in **Table 1**.

**BOX 1.**
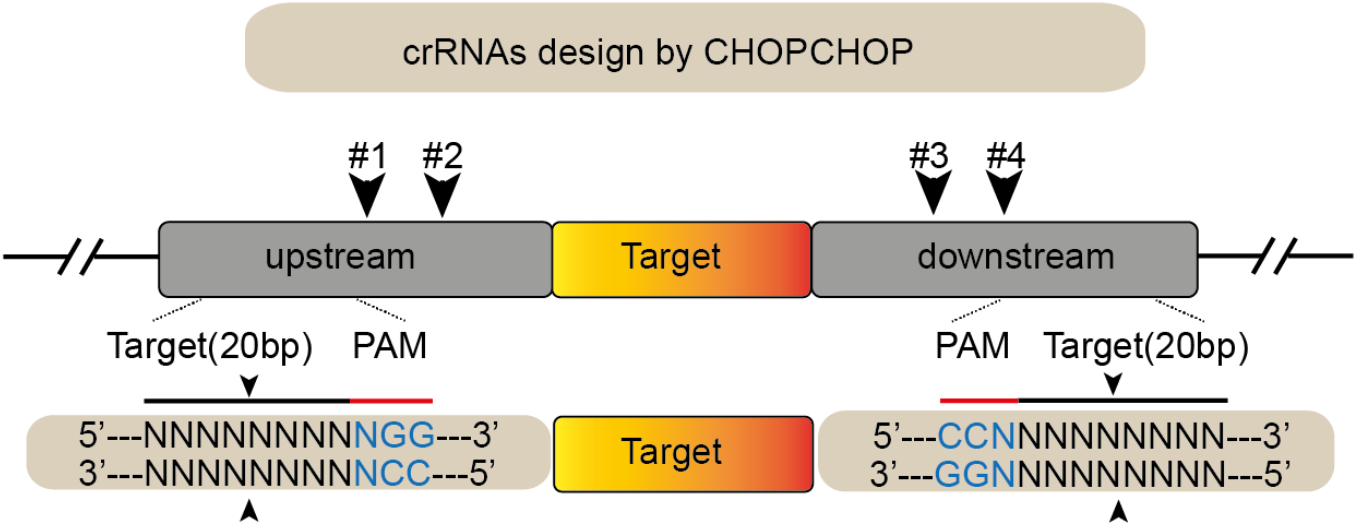
Schematic of CRISPR RNAs. Designing crRNAs for a defined target locus is critical for the successful implementation of the MASTR-seq protocol. We design 4 distinct crRNAs specific to upstream and downstream flanking sequences of a target STR using the CHOPCHOP tool.

#### Samples

MASTR-seq is illustrated here using data generated in patient-derived human induced pluripotent stem cells (hiPSCs) and human embryonic stem cells (hESCs), but it is adaptable to diverse cell types or tissues. The culture medium formulation, seeding density, growth and passage conditions, and cell or tissue dissociation methods will be specific to the cell type under investigation.

#### Cell number

We used TrypLE reagents to triturate hiPSCs into a single-cell suspension. We counted single iPSCs using an automated Countess system or manual counting with a hemocytometer. We start with 5-10 x 10^6 cells for each sample as input to achieve a minimum of 30 μg of high molecular weight (HMW) genomic DNA, which is sufficiently used for 5x Cas9 RNP complex cleavages. Future versions might be optimized to allow for low input DNA from rare samples.

**BOX 2.**
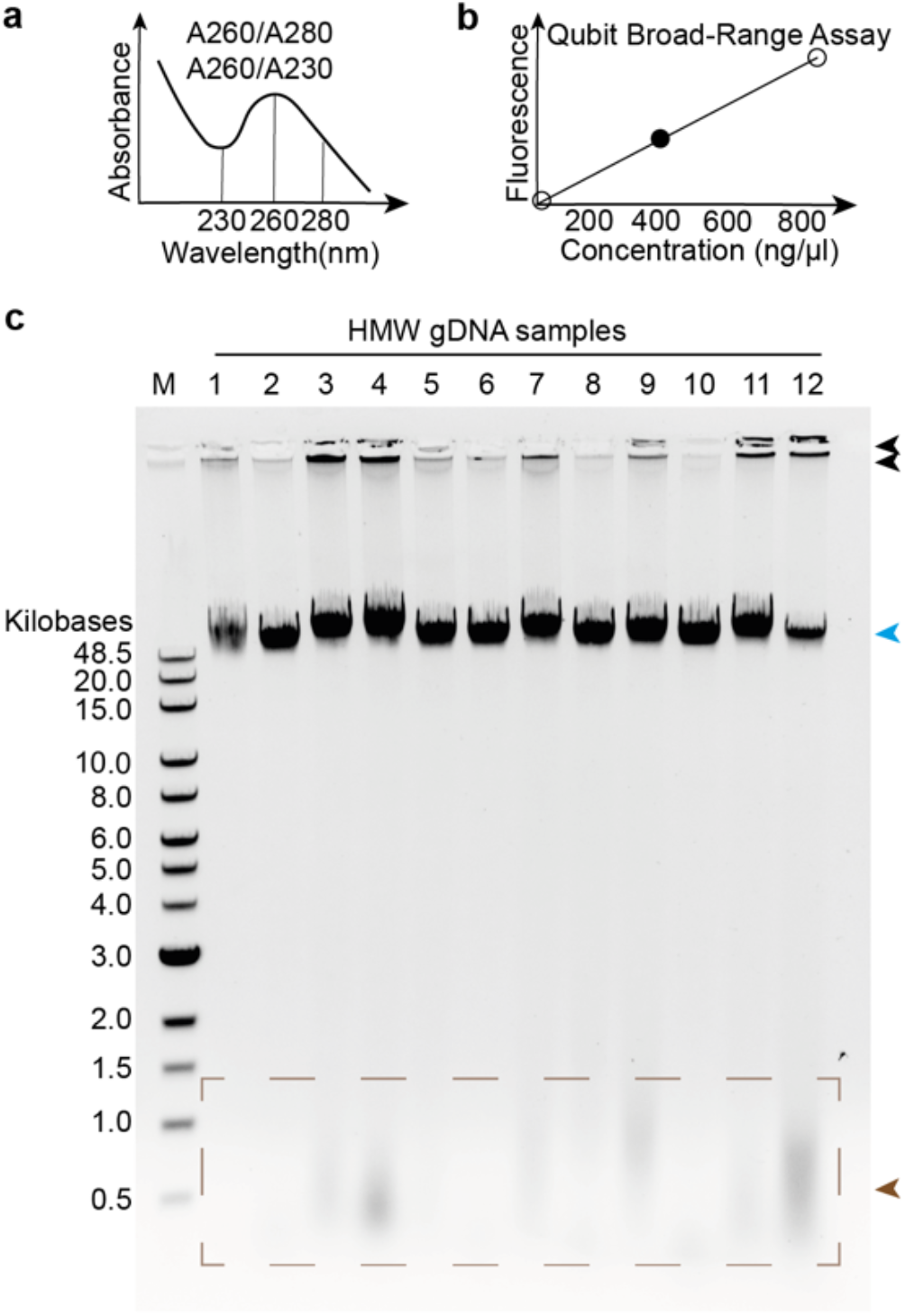
Quality control assays for high molecular weight genomic DNA. Using high-molecular weight (HMW) genomic DNA of high quality is critical to successful MASTR-seq data generation. (**a**) We measured the purity (A260/A280 and A260/A230 ratios) of gDNA with Nanodrop. (**b**) We measured the concentration of rehydrated genomic DNA using the Qubit Broad-Range assay. (**c**) We checked the integrity of HMW gDNA by running samples on a low concentration (0.6%) TAE agarose gel. Black and blue arrows mark HMW genomic DNA. Degraded DNA fragments can be visualized at the bottom of gels marked by brown arrowheads.

#### High molecular weight (HMW) genomic DNA preparation

DNA integrity is essential for efficient yield of Cas9 target cleavage products. Therefore, we employ the phenol–chloroform extraction method to extract HMW DNA exceeding 100 kb in length. DNA extracted using kits often results in fragments ranging from 10–100 kb in length^19^. Moreover, HMW DNA tends to be viscous, and the rehydration process can lead to fragmentation. After extraction, we recommend eluting HMW genomic DNA by heating at 50°C for 1 hour, followed by rotation at room temperature overnight. We measure genomic DNA yield as the concentration of rehydrated DNA using the Qubit DNA Broad-Range assay. Genomic DNA purity is assessed by calculating the A260/A280 and A260/A230 ratios obtained through spectrophotometric analysis with Nanodrop. High purity genomic DNA should exhibit A260/A280 and A260/A230 ratios within the ranges of 1.8-2.0 and 2.0-2.2, respectively. If the ratios significantly deviate from these optimal values, superior performance can be achieved by re-purifying gDNA samples using AMPure® XP reagent. We verify the integrity of HMW DNA fragments on a 0.6% 1X TAE agarose gel (**BOX 2**). Using iPSC pellets consisting of 5-10 x 10^6 cells for each sample, we typically observe a total mass of 30-60 μg of high molecular weight (HMW) genomic DNA. Alternatively, some Cas9-targeted long-read methods employ genomic kits, such as the Gentra Puregene Cell Kit (Qiagen, 158767) for HMW DNA extraction, which also yields satisfactory results. However, in cases with limited samples, the phenol–chloroform extraction method may be preferred to maximize genomic DNA yield.

#### Cas9 ribonucleoprotein complex (RNP) assembly

The CRISPR-Cas9 system configured as a ribonucleoprotein (RNP) complex *in vitro* consists of a guide RNA and Cas9 nuclease. The guide RNA is comprised of two components: (1) a 17-20 nucleotide crRNA complementary to the target DNA and (2) a tracrRNA which acts as a binding scaffold for the Cas9 nuclease. To assemble Cas9 RNP complexes, we anneal crRNAs and tracrRNA in a 1:1 molar ratio. For example, when targeting the *FMR1* gene for 10 reactions, we combine 4 different crRNAs, each pooled with tracrRNA (2.5 μM for each crRNA and 10 μM for tracrRNA in a 10 μL reaction volume). Next, we assemble CRISPR-Cas9 RNPs in 1X NEB CutSmart buffer. We determine the optimal ratio of Cas9 protein to CRISPR gRNA based on *in vitro* Cas9 RNPs cleavage efficiency. In this protocol, we employed a 1:1 molar ratio of Cas9 protein to CRISPR gRNA to achieve efficient *FMR1* target cleavage (**BOX 3**).

**BOX 3.**
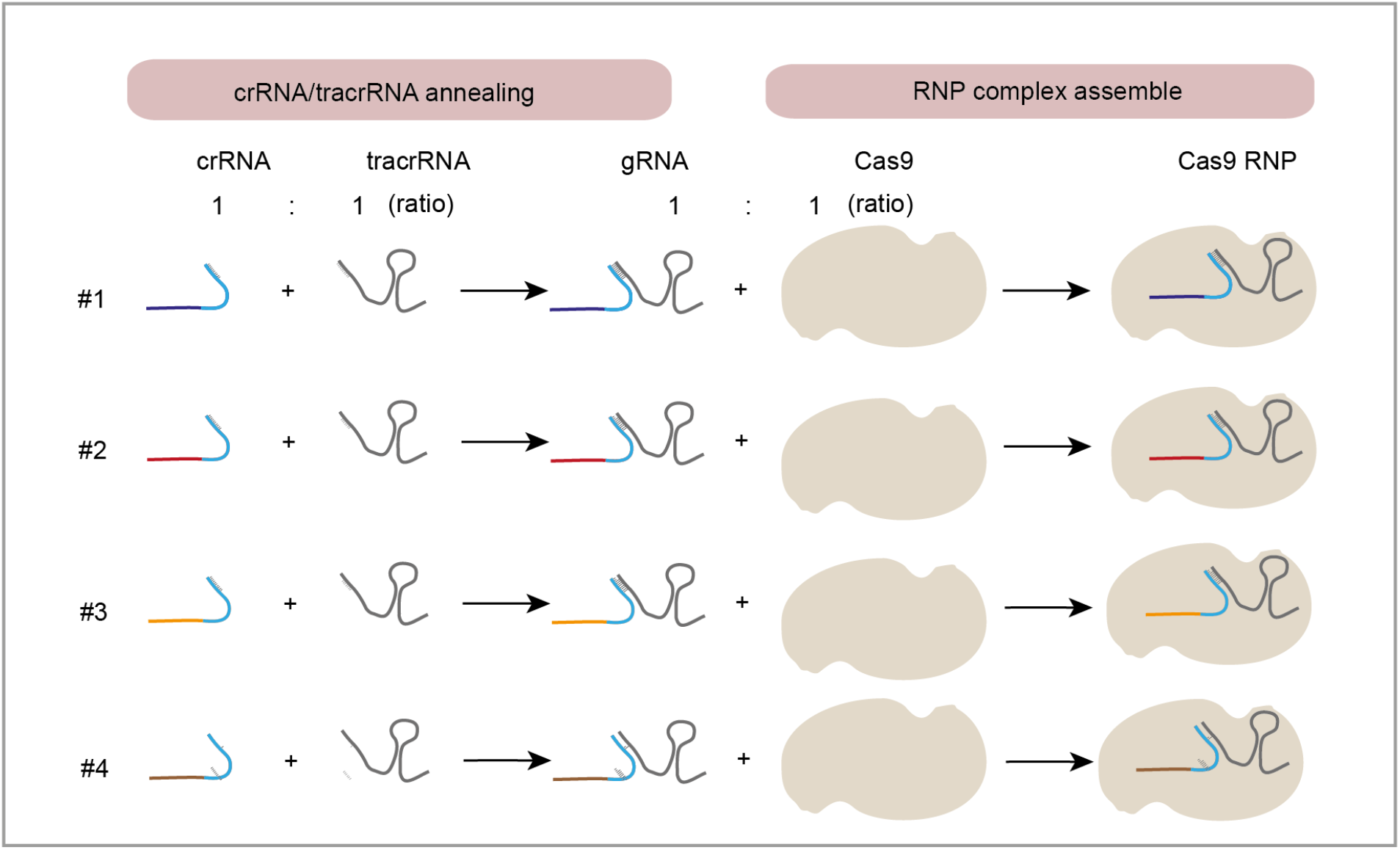
Optimization of Cas9 RNP assembly. To determine the optimal ratio of Cas9 protein to CRISPR guide RNA, we measured Cas9 RNP cleavage efficiency *in vitro*. Here, we use a 1:1 molar ratio of Cas9 protein to each distinct CRISPR gRNA, resulting in efficient *FMR1* target cleavage.

#### HMW genomic DNA dephosphorylation

Genomic DNA dephosphorylation is the initial step to reduce genomic background in MASTR-seq (Figure 1, step 3). We incubate 5 μg of genomic DNA with 3 μL Quick CIP for DNA dephosphorylation in NEB rCutSmart Buffer at 37°C for 20 minutes followed by inactivation at 80°C for 2 minutes. When optimizing for increased on-target coverage, it is recommended to divide genomic DNA into multiple reactions in the dephosphorylation step, each containing 5 μg (or less) per reaction.

#### Cas9 RNP cleavage and dA tailing

We incubated Cas9 RNPs with a maximum of 5 μg of dephosphorylated HMW genomic DNA at 37°C for 60 minutes followed by a dA-tailing of blunt ends at 72°C for 5 minutes. For efficient on-target cleavage, we recommend an optimization of Cas9 RNP cleavage time to increase the ratio of on-target/off-target reads. Excessive cleavage time can lead to a higher number of off-target reads. To measure Cas9 RNP cleavage efficiency, we recommend designing primers flanking the Cas9 RNP cleavage locus and performing PCR to amplify the CRISPR-targeted locus (**BOX 4**).

**BOX 4.**
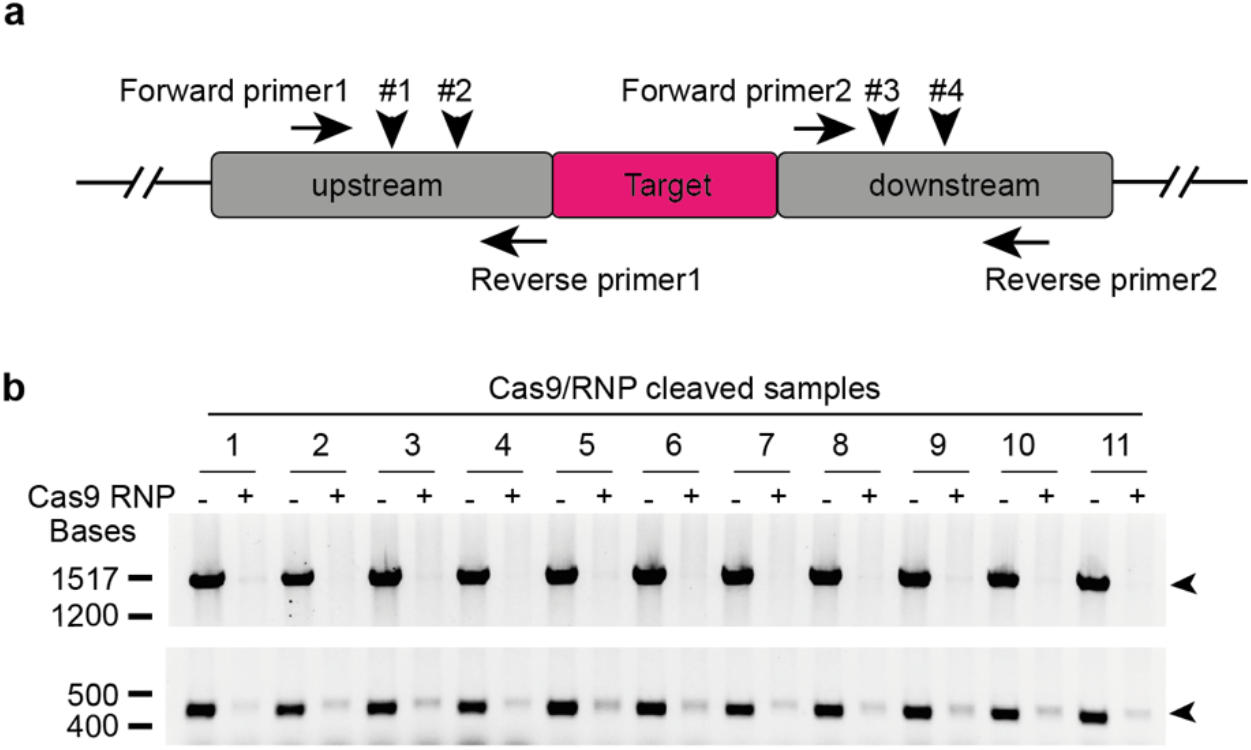
Cas9 RNP cleavage efficiency assay. Accurately assessing the efficiency of Cas9 RNP cleavage is crucial for optimizing the on-target read percentage of MASTR-seq. (**a**) To assess the efficiency of Cas9 RNP cleavage, we have designed two pair primers that flank the Cas9 RNP cleavage site at upstream and downstream of target region, respectively. (**b**) Upon PCR amplification of the Cas9-targeted locus, compared to negative controls (-), we observe that signals marked by arrowheads from samples with Cas9 RNP cleavage (+) is nearly completely/mostly lost. The top arrowhead indicates PCR amplification product of upstream region using Forward primer1/Reverse primer1 shown in (**a**), while the bottom arrowhead indicates PCR amplification product of downstream region using Forward primer2/Reverse primer2 shown in (**a**).

#### DNA purification

We employed the sodium acetate-ethanol DNA precipitation method to purify and concentrate DNA from high-salt aqueous solutions. The precipitated DNA is rehydrated at 50°C for 1 hour in 30 μL elution buffer. If DNA is not completely dissolved, an overnight rotation at room temperature is recommended.

#### BluePippin automated size selection

Removing background uncleaved genomic DNA or off-target DNA fragments outside of the target locus is essential for MASTR-seq performance. We use BluePippin size selection to further enrich the target signal in our protocol. Cas9 RNP cleavage of the target (e.g. the *FMR1* locus) produces a 6.9 kb-size DNA fragment in normal-length iPSCs or a 8-10 kb sized DNA fragments in mutation-length iPSCs. We conduct size selection for large 7-10 kb fragments using the BluePippin (Sage Science) and BLF7510 cassette using the ‘0.75DF 3-10 kb Marker S1-Improved Recovery’ cassette definition with broad range mode set at 5-12 kb (**Table 2**). We recommend testing different size range modes before finalizing the size selection. To allow for size selected HMW DNA to efficiently detach from the membrane of elution mode well, we recommend allowing samples to remain in the cassette for at least 45 mins after the run is complete prior to removing eluent. Additionally increased yield can be achieved by second elution with 0.1% Tween 20 solution after removal of initial elution.

**BOX 5.**
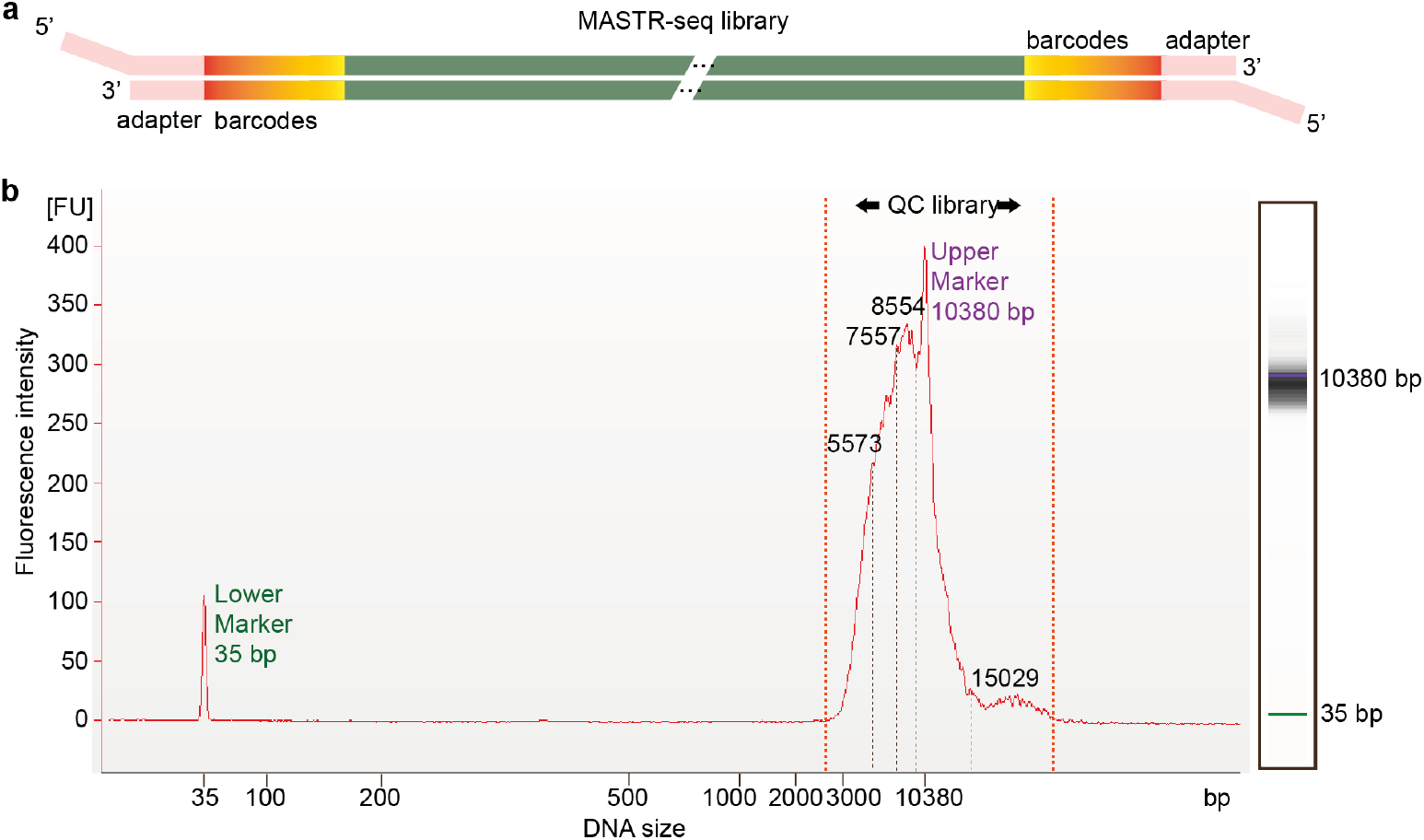
Bioanalyzer profile of a final MASTR-seq library. When performing MASTR-seq for the first time, we recommend assessing the quality of the library by performing a high sensitivity DNA assay on a 2100 bioanalyzer or Tape Station. **(a)** The successful library should contain Cas9 RNP cleaved DNA fragments with both adaptor and unique barcode addition. We have used a 2100 bioanalyzer to analyze the concentration and size distribution of a long-read library. **(b)** The resulting electropherogram shows size selected broad peaks corresponding to the DNA fragments ranging from 3 kb to 15 kb, which reflects Bluepippin target size range (5-12 kb).

#### PCR-free multiplexed barcoding

To prepare the library for nanopore sequencing, we ligate size selected DNA fragments to a unique barcode for each sample, pool multiple barcoded samples together, and ligate to an adapter for sequencing. To circumvent amplification bias over repetitive sequences, our methods are PCR free. The limitation is that the low yield of target DNA fragments makes equimolar pooling of each barcoded sample challenging. Using MASTR-seq, we can barcode each sample and pool 8-12 multiplexed pooled human derived samples for one MinION flow cell. Library quality can be assessed by using the Agilent bioanalyzer or Tape station system (**BOX 5**). The MinION flow cell is limited by 512 nanopore channels for sequencing and the PromethION flow cell contains up to 2675 nanopore channels. For applications using the PromethION flow cell, MASTR-seq can be readily adapted to increase sample throughput and the number of target loci. In principle, one PromethION flow cell can accommodate over 50 multiplexed samples.

#### Sequencing

To enhance the yield of on-target reads and reduce off-target reads during sequencing, we employed specific parameters such as a minimal read length of 1000 bp, a maximum scanning time of 1.5 hours, and “pores reserve” mode (**Table 3**). For sequencing, we typically utilize the Flow Cell R9.4.1 on the MinION sequencer for a duration of 48 hours to achieve more than 95% yield. To further improve the yield and accuracy of long reads, it is advisable to implement MASTR-seq with the upgraded R10.4.1 flow cell in conjunction with the more precise Q20+ chemistry.

#### Data processing

##### a. Raw data processing

We conducted base-calling of raw data (fast5 files) using guppy_basecaller (version 6.2.1). Following base-calling which generated sequencing summary file (**Table 4**), we assessed data yield across sequencing time (**BOX 6a**) and pass/fail reads quality distribution (**BOX 6b**). We then used guppy_barcoder (version 6.2.1) to demultiplexed base-called reads into unique sample and generated barcoding summary file (**Table 5**) and analyzed the distribution of reads quality across each de-multiplexed sample (**BOX 6c**). We generated a summary of on-target reads data using the Cas9 enrichment workflow from EPI2ME Labs application (**Table 6**). We aligned demultiplexed reads to the hg38 human reference genome using minimap2 (version 2.22-r1101). Finally, we computed the on-target coverage per sample using ‘Samtools’ (**BOX 6d**). We demonstrated that MASTR-seq has >10-fold more proportion of on-target reads than the previously published method nCATS^33^ (**BOX 6e, Table 7**).

**BOX 6.**
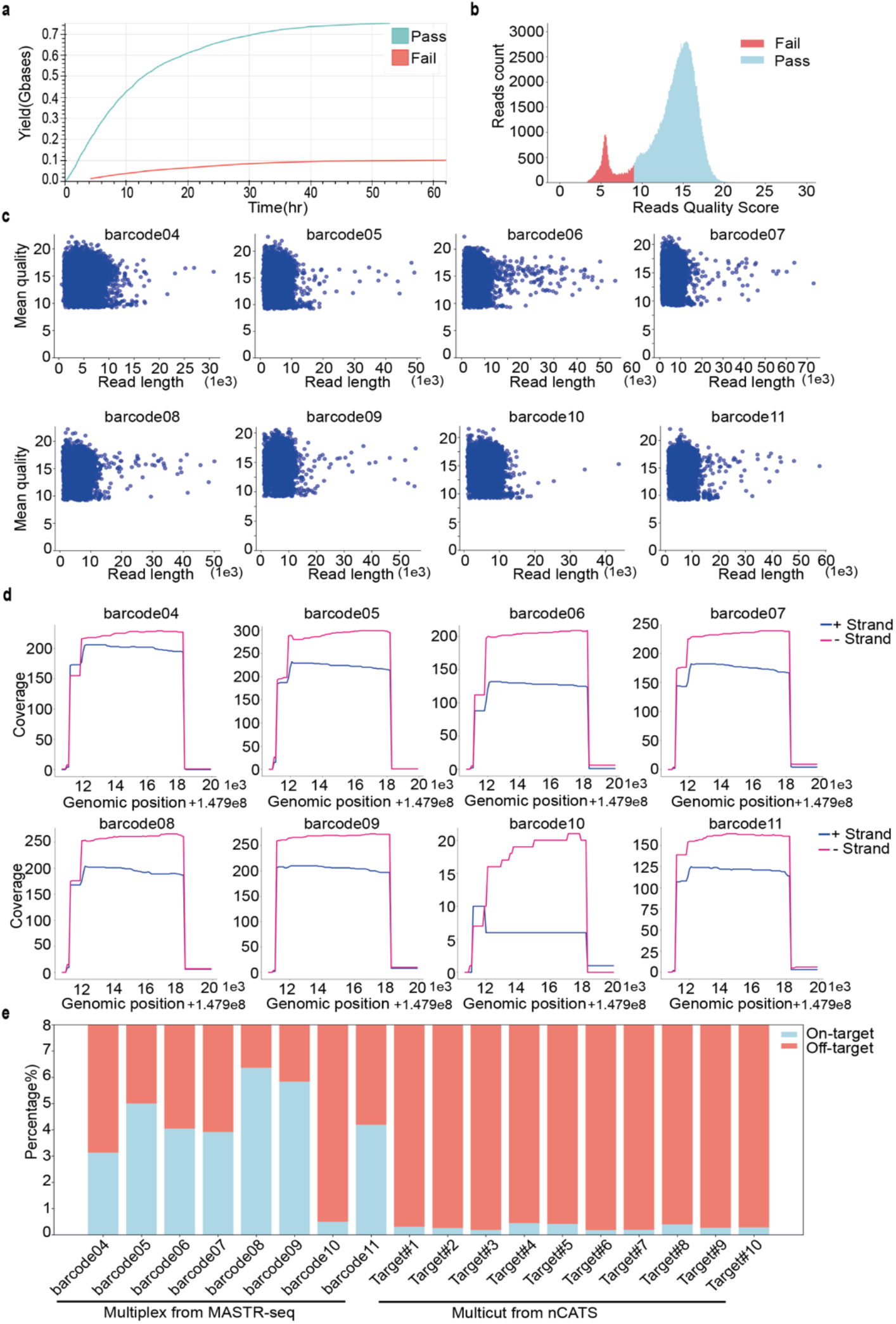
Sequencing summary of *FMR1* CGG STR target across multiple indexed samples by MASTR-seq. To assess the yield and quality of long reads, we conducted an analysis of one nanopore run(run_id,FAS91717). (**a**) A representative summary of sequencing shows the percentage of pass (blue) and fail (red) reads. (**b**) The distribution of read quality score for a representative MASTR-seq library. quality and on-target coverage of MASTR-seq data (**c**) Distribution of read quality score versus read length across multiple indexed samples. (**d**) On-target read coverage stratified by sense and antisense strands across multiple indexed samples. (**e**) Comparation of the proportion of on-target versus off-target reads for 8 multiplexed samples (from barcode04 to barcode11, Table 7, Run_id FAS91717) multiplexed in one MASTR-seq library and the previously published nCATS library data (from target#1 to target#10, Table 7, Run_id TG_09)^33^.

In the future, data processing improvements might lead to improved base-calling or read alignment. Several studies report that higher accuracy can be achieved by using R10.4 flow cells^57-59^. R10.4 offers improved resolution for homopolymer detection compared to R9.4.1^57, 58^. Using R10.4 flow cell, the simplex sequencing mode demonstrates an average read quality of 98%, while duplex reads exhibit 99% accuracy^58^. Moreover, by implementing the latest base-calling algorithm (dorado) for processing data produced from the flow cell (R10.4.1) together with Q20+ reagent, this combination claims an overall 99% reads accuracy^60, 61^.

##### b. STR visualization

Strand-specific errors in nanopore base calling are common in repetitive DNA sequences^62^. When assessing STR tract length, we compared the forward with reverse strands. We found that only the reverse reads accurately reflected the length of the *FMR1* CGG STR, whereas both forward and reverse reads measured similar *HTT* CAG STR lengths (see **Figures 2-4**). We utilized customized scripts for the visualization of STRs. To ensure the use of only high-quality reads in downstream analysis of *FMR1* CGG STR, we implemented several quality control steps: (1) removing reads that do not align to the target region and (2) using only reads that mapped to the reverse strand which has less errors. For the *FMR1* locus, we filtered out truncated reads lacking an upstream sequence to the CGG tract “ACCAAACCAA” and keep reads with at least four consecutive CGGs. We also excluded reads containing more than nine consecutive “TA” nucleotides within the CGG STR, which we find is indicative of base calling errors. For the *HTT* locus, we removed reads that do not align to the target region, used reads that map to the reverse or forward strand, and kept reads with at least three consecutive CAGs. Finally, we quantified the number of repeat units in the continuous STR tract and plotted representative nanopore long-reads for visualization (see **Figures 2-4**). To streamline data processing for STR visualization, we developed a simplified python tool named MASTRseq.py (see bitbucket links provided below). The tool processes the FASTQ and SAM files as input, produces the STR counts file, and creates plots to illustrate the distribution of STRs across reads for each sample.

**Figure 2.**
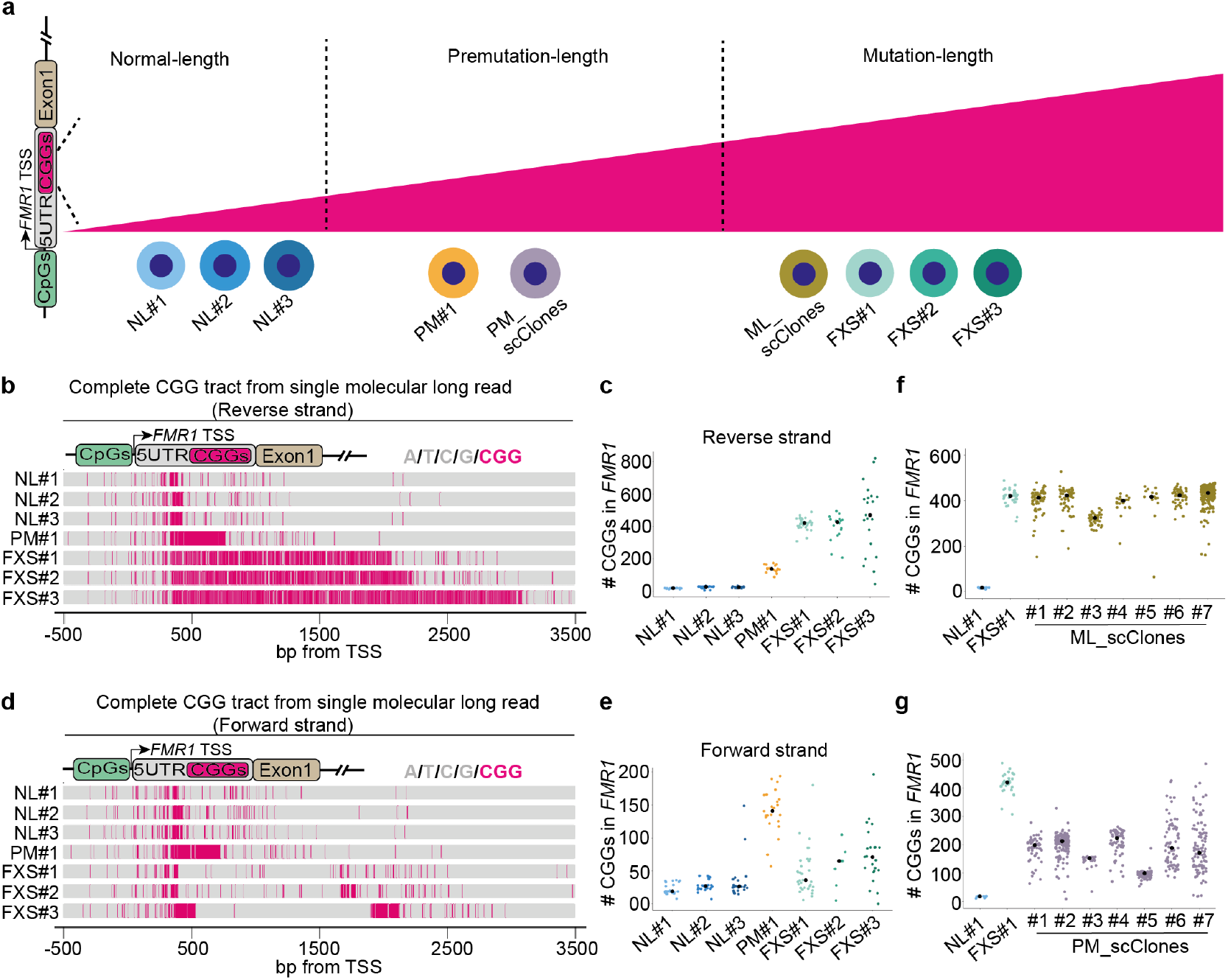
Precise quantification of *FMR1* CGG STR lengths in human iPSCs derived from fragile X syndrome (FXS) patients using MASTR-seq. **(a)** Schematic representation of iPSC lines modeling normal-length (NL), premutation-length (PM), and mutation-length (ML/FXS) *FMR1* CGG STR tracts. **(b)** Representative reverse nanopore long-read spanning the 5’ UTR of *FMR1*. Nucleotides are color-coded (gray: A/T/C/G, Pink: CGG). **(c)** Distribution of CGG STR tract length in *FMR1* 5’ UTR derived from individual forward nanopore long reads. Each dot represents a single molecule long read, corresponding to one allele. **(d)** Representative forward nanopore long-read along the *FMR1* 5’ UTR. Nucleotides are color-coded (gray: A/T/C/G, Pink: CGG). **(e)** Distribution of CGG STRs in the 5’ UTR of *FMR1* obtained from individual reverse nanopore long-read. Each dot signifies a single molecule long read, corresponding to one allele. **(f-g)** Number of CGG STRs in the 5’UTR of *FMR1* computed from individual reverse single-molecule nanopore long reads across single-cell iPSC clones derived from **(f)** a FXS mutation-length line and **(g)** a premutation-length line. These data are provided side by side with the population-based mutation-length (FXS#1) and normal-length (NL#1) lines.

#### DNA methylation

We employed distinct methods for calling DNA methylation at the *FMR1* promoter region (500 bp upstream to TSS) and *FMR1* CGG tract, respectively. We utilized nanopolish (version 0.13.2) to identify CpG methylation within the 500 bp *FMR1* promoter (hg38, chrX: 147911419-147911919). Given nanopolish’s limitation in calling DNA methylation over a variable number of CGG triplets, we used STRique (version 0.4.2) to call methylation of CpGs over the CGG tract in normal-length, premutation-length, and mutation-length iPSCs.

For the quantification of CpG methylation at the *FMR1* or *HTT* promoter using nanopolish, we indexed the fast5 files using the ‘Index’ command. We applied the ‘call-methylation’ command to detect CpG methylation within the specified windows of GRCh38-hg38 for *FMR1* loci (chrX:147,902,117-147,960,927) and *HTT* loci (chr4:3,072,146-3,078,241). Methylated and unmethylated states were computed based on log2 likelihood values, with >0.1 considered as methylated. The proportion of methylated CpGs is computed for each single molecule read. The resulting proportions are visualized as kernel distribution estimation plots using the ‘density’ function in R (see **Figures 3**).

**Figure 3.**
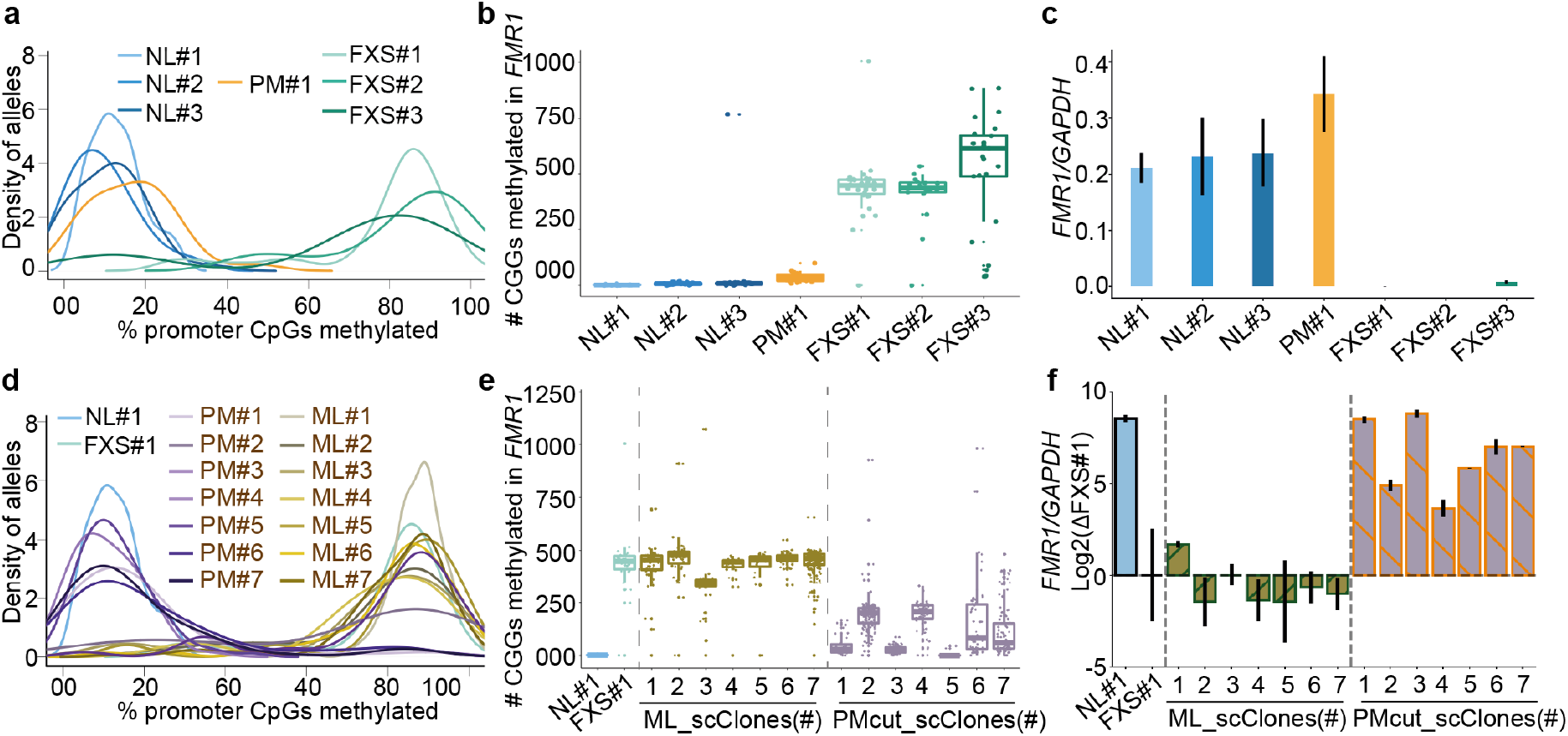
Precise quantification of DNA methylation and STR tract length in the same single molecule in iPSCs derived from FXS patients using MASTR-seq. **(a)** CpG dinucleotide DNA methylation of the *FMR1* promoter across normal-length (NL), premutation-length (PM), and full mutation-length (FXS/ML) iPSC lines. Histogram of methylated CpG islands in the *FMR1* promoter from reverse nanopore long-reads computed using nanopolish. **(b)** Proportion of alleles with a methylated CGG STR tract using reverse nanopore long reads computed using STRique across NL, PM, and ML iPSC lines. Each dot denotes a single molecule long read representing one allele. **(c)** qRT-PCR measurement of *FMR1* mRNA levels normalized to *GAPDH* mRNA levels. Horizontal bar indicates the mean value from 2 biological replicates. Error bars represent the standard deviation between 2 biological replicates. **(d-e)** CpG dinucleotide DNA methylation of the **(d)** *FMR1* promoter and **(e)** CGG STR tract computed from individual reverse single-molecule nanopore long reads across 7 single-cell iPSC clones derived from a FXS ML line and 7 premutation-length CRISPR cutback clones also derived from the same parent FXS ML iPSC line. These data are provided side by side with the population-based ML (FXS#1) and NL (NL#1) iPSC lines. **(f)** qRT-PCR measurement of *FMR1* mRNA levels normalized to *GAPDH* mRNA levels for each single-cell clone and further normalized to the values for parent line FXS#1. Horizontal bar indicates the mean value from 2 biological replicates. Error bars represent the standard deviation between 2 biological replicates.

To ascertain CpG methylation specifically at the CGG STR in the 5’UTR of *FMR1* using STRique, we processed the fast5 files with the ‘index’ command. We computed methylation status and CGG counts using the ‘count’ command with the respective models ‘r9_4_450bps_mCpG.model’ and ‘r9_4_450bps.model’. We retained reads with prefix and suffix scores greater than 4 for further analyses. Those with scores <4 exhibit low-quality mapping to the upstream and downstream regions of the CGG tract. Subsequently, we calculated the number of methylated CGGs and presented them as jitter plots using the R ggplot2 package (see **Figures 3**).

#### Recommended controls in the procedures

To evaluate the efficiency of Cas9 RNP cleavage, we recommend preparing the negative control samples (non Cas9 RNP cleavage) as described in Day 3. To evaluate the efficiency of Cas9 RNP cleavage at new targets, we recommend performing a positive control group. We recommend using cell lines with defined STR tract lengths to calibrate the method’s parameters to ensure MASTR-seq is accurately measuring ground-state truth before applying it to query unknown cell lines (see **Figures 4**).

**Figure 4.**
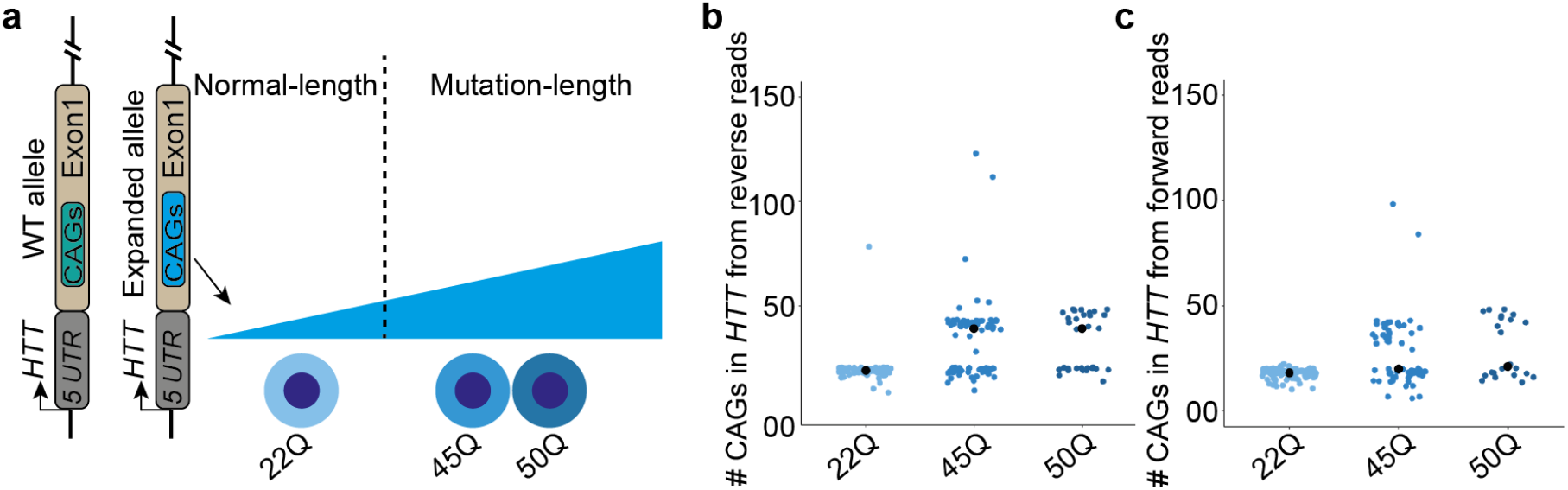
Precise quantification of *HTT* CAG STR tract lengths using MASTR-seq. **(a)** Schematic representation of isogenic REUS2 human embryonic stem cell (hESC) lines engineered to model *HTT* CAG expansion in Huntington’s disease (HD). Shown here are REUS2 subclones comprising one *HTT* CAG normal-length and two *HTT* CAG mutation-length. **(b, c)** Distribution of CAG triplets in *HTT* obtained from individual **(b)** reverse and **(c)** forward strand nanopore long reads. Each dot signifies a single molecule long read corresponding to one allele.

#### Biological materials

Induced pluripotent stem cell (iPSC)

Normal-length (NL), premutation-length (PM), and full mutation-length (ML) FXS-derived human iPSCs published in our recent public work^48^ (**Figure 2a**). *HTT* CAG normal-length and disease-length hESC pellets were acquired from Dr. Kristen Brennand, and the hESC lines were created by Brivanlou lab at Rockefeller University, NY^63^ (**Figure 4a**).

**Reagents**

mTeSR Plus media (STEMCELL Technology, cat. no. 100-0276, 100-1130)

Matrigel hESC-Qualified Matrix (Corning, cat. no. 354277)

DMEM/F-12 (Gibco, cat. no. 11320033)

1X PBS without calcium and magnesium (Corning, cat. no. 21-040-CV)

Penicillin-streptomycin (Gibco, cat. no. 15140122)

RevitaCell™ Supplement (100X) (Gibco, cat. no. A2644501)

Synth-a-Freeze (Gibco, cat. no. A1254201)

TrypLE™ Express Enzyme (1X), phenol red (Gibco, cat. no. 12605010)

Versene Solution (Gibco, cat. no. 15040066)

10X TAE buffer (Invitrogen, cat. no. 15558042)

Agarose (Lonza, cat. no. 50004)

1M Tris-HCl (pH 8.0) (Invitrogen, cat. no. 15568025)

1X TE Buffer (pH 7.5) (Integrated DNA Technologies, cat. no. 11-05-01-15)

0.5M EDTA (pH 8.0) (Invitrogen, cat. no. 15575020)

20% SDS (w/v) solution (Fisher Scientific, cat. no. BP1311-200)

RNase A (Sigma-Aldrich, cat. no. 10109142001)

Proteinase K (New England Biolabs, cat. no. P8107S)

UltraPure Phenol/Chloroform/Isoamyl Alcohol (Invitrogen 15593049)

5M Ammonium Acetate (Invitrogen, cat. no. AM9070G)

100% Ethanol (Decon Labs, cat. no. 2716)

0.75% Agarose Gel Cassette (Sage Science, cat. no. BLF7510)

Ligation Sequencing Kit (Oxford Nanopore Technologies, cat. no. SQK-LSK109)

Native Barcoding Expansion 1-12 (Oxford Nanopore Technologies, cat. no. EXP-NBD104)

Flow Cell Priming Kit (Oxford Nanopore Technologies, cat. no. EXP-FLP002)

S. pyogenes Cas9 tracrRNA (Integrated DNA Technologies, cat. no. 1072532)

Alt-R® S. pyogenes HiFi Cas9 nuclease V3 (Integrated DNA Technologies, cat. no. 1081060)

Quick Calf Intestinal Phosphatase (New England Biolabs, cat. no. M0525S)

rCutSmart buffer (New England Biolabs, cat. no. B6004S)

Nuclease-free water (ThermoFisher, cat. no. AM9937)

Taq DNA polymerase (New England Biolabs, cat. no. M0273S)

100 mM dATP solution (Thermo Scientific, cat. no. R0141)

Nuclease-free duplex buffer (Integrated DNA Technologies, cat. no. 11-01-03-01)

Quick-Load^®^ 1 kb Extend DNA Ladder (New England Biolabs, cat. no. N3239S)

NEB Blunt/TA Ligase Master Mix (New England Biolabs, cat. no. M0367L)

T4 DNA ligase, from NEBNext Quick Ligation Module (New England Biolabs, cat. no. E6056S)

Agencourt AMPure XP beads (Beckman Coulter, cat. no. A63881)

Qubit™ Broad Range dsDNA Quantitation (Invitrogen, cat. no. Q32850)

Qubit™ High Sensitivity dsDNA Quantitation (Invitrogen, cat. no. Q32851)

Bioanalyzer High Sensitivity DNA Analysis Kit (Agilent, cat. no. 5067-4626)

#### Equipment

Cell culture incubator (Thermo Scientific, cat. no. 51026332)

R9.4.1 flow cell (Oxford Nanopore Technologies, FLO-MIN106D)

Countess 3 automated cell counter (Thermo Scientific, Cat. no. AMQAX2000)

Hemocytometer (Fisher Scientific, cat.no. 0267151B)

BluePippin (Sage Science)

Agilent bioanalyzer (Agilent Technologies, model no. G2939A)

MinION Mk1B Sequencing Device (Oxford Nanopore Technologies, MIN-101B)

Qubit™ 2.0 Fluorometer (Invitrogen, cat. no. Q32866)

Nanodrop 2000 Spectrophotometer (Thermo Scientific, cat. no. ND-2000)

Owl™ EasyCast™ B2 Mini Gel Electrophoresis device (Thermo Scientific, Cat. no. 09-528-118)

ChemiDoc™ MP Imaging System (Bio-rad,cat.no. 12003154)

Magnetic rack (DynaMag-2 Thermo Fisher Scientific, cat nos. 12321D)

Vortex-Genie® 2 mixer (Sigma-Aldrich, cat. no. Z258415)

Tube Revolver Rotator (Thermo Scientific, cat. no. 88881001)

Applied Biosystems™ 96 Well Thermal cycler (Thermo Scientific, cat.no. 4375305)

ART™ Wide Bore Filtered Pipette Tips (Thermo Scientific, cat. no.2069G)

MaXtract High Density 50 mL tubes (Qiagen, cat. no. 129073)

50mL Centrifuge Tube (VWR, cat. no. 89039-656)

2.0 mL DNA LoBind® Tubes (Eppendorf cat. no. 022431048)

1.5 mL DNA LoBind® Tubes (Eppendorf cat. no. 022431021)

0.2 mL PCR 8-Tube Strips (USA Scientific, cat.no. 1402-4700)

#### Computational sources (software and algorithms)

PC used to plug in MinION Mk1B Sequencer

High-performance computing cluster (optional)

ChopChop online tool version 3.0.0^64^ https://chopchop.cbu.uib.no/

MinKNOW version 22.08.9 Oxford Nanopore Technologies, https://community.nanoporetech.com/downloads

Nanopolish (version 0.14.0)^65^ https://github.com/jts/nanopolish

Minimap2 (version 2.22-r1101)^66^ https://github.com/lh3/minimap2

Samtools (version 1.11)^67^ https://www.htslib.org

Guppy (version 6.2.1) Oxford Nanopore Technologies. https://community.nanoporetech.com/downloads

STRique (version 0.4.2)^28^ https://github.com/giesselmann/STRique

EPI2ME (version 5.1.9) Oxford Nanopore Technologies. https://labs.epi2me.io

#### Reagent setup

**⚠ CRITICAL**

We recommend storing all buffers at 4 °C unless otherwise stated in the protocol.

#### Human iPS cells culture medium

Prepare mTeSR Plus media by mixing 400 ml mTeSR Plus basal medium with 100 mL 5X supplement, supplemented with 1% (v/v) penicillin-streptomycin. Store media at 4 °C and bring to room temperature (RT) before use.

#### Cell dissociation buffer

Prepare 0.5X TrypLE cell dissociation buffer at RT by diluting 1 volume of 1X TrypLE with 1 volume of 1X PBS without calcium and magnesium.

#### 10 mM Tris-HCl (pH 8.0) buffer

Prepare 10 mM Tris-HCl (pH 8.0) buffer by adding 0.1 mL 1 M Tris-HCl (pH 8.0) to 9 mL nuclease-free water at RT.

#### crRNA stock solution (100 μM)

Prepare 100 μM crRNA stock solution by resuspending 5 nmol crRNA in 50 μL TE (pH 7.5). Store aliquots at -80 °C and thaw on ice before use.

#### tracrRNA stock solution (100 μM)

Prepare 100 μM tracrRNA stock solution by resuspending 5 nmol tracrRNA in 50 μL TE (pH 7.5). Store aliquots at -80 °C and thaw on ice before use.

#### Cell lysis buffer

Prepare 50 mL of fresh cell lysis buffer at RT by mixing 500 μL of 1 M Tris-HCl (pH 8.0), 400 μL of 0.5 M EDTA (pH 8.0), 1250 μL of 20% SDS (w/v) in nuclease-free water.

#### Preparation of the Matrigel-coated plate

Coat iPSC culture plates with 1.2% (v/v) Matrigel hESC-Qualified Matrix in DMEM/F-12 for at least 1 hour at 37 °C.

#### Procedure

Note: Perform all below steps at room temperature are 24 °C unless otherwise stated in the protocol.

#### Cell culture and cell counting

1. Culture human iPSC lines in mTeSR Plus media supplemented with 1% (v/v) penicillin-streptomycin on Matrigel hESC-Qualified Matrix-coated plates at 37 °C and 5% CO_2_. To maintain the physical separation of single colonies without merging, routinely passage cells when they reach 60-70% confluence every 4-5 days. For iPSC passaging, incubate plates in Versene solution at 37 °C for 5 minutes, followed by deactivation of Versene with 2X volume of mTeSR Plus media, spin down cell pellets at 200 xg for 3 minutes. Resuspend hiPSC pellets with mTeSR Plus media and split cell suspension into multiple new Matrigel matrix-coated plates.

#### ⚠ CAUTION

To eliminate potential STR expansion/contraction during cell passages, all iPSCs should be expanded and cryopreserved at low passage numbers.
2. To count cells, incubate hiPSCs with 0.5X TrypLE cell dissociation buffer on the plate for 4 mins. Then gently pipette to dissociate human iPSC colonies into single cells.
3. Take 100 μL of suspended cells into a new microcentrifuge tube and add 100 μL 0.4% Trypan Blue (final concentration 0.2%). Mix gently and incubate for 2 mins at RT.
4. Count cell numbers manually with a hemocytometer or use the Countess automated cell counter to get cell numbers.
5. Aliquot 5-10 million cells into each 2 mL microcentrifuge tube, spin down at 300 xg for 5 mins to pellet the cells, remove supernatant, and flash-freeze pellets in liquid nitrogen. Store pellets at -80 °C until use.

#### HMW genomic DNA preparation and quality assay

6. Prepare a cell lysis buffer (1.0 mL per 1 million cells) and add RNase A at a final concentration of 20 μg/mL.
7. Resuspend 5 million iPSC pellets in 100 μL of PBS, add 4.9 ml lysis buffer with RNase A included, and mix gently with a wide bore pipette tip. Incubate for 1 hour at 37 °C.

##### ⚠ CAUTION

To ensure thorough cell lysis, incubate samples at 37 °C for 30 minutes, and then gently pipette using wide-bore tips to mix and continue incubation for 1 hour at 37 °C.

##### Ⓘ TROUBLESHOOTING

8. Add proteinase K to a final concentration of 4 units/mL. Gently invert to mix, and then incubate at 50 °C for 3 hours. During this process, centrifuge the MaXtract High Density tube at 4000 xg for 2 minutes to spin down the gel pellet.
9. Pour each lysed sample evenly into MaXtract High Density 50 mL tubes. Add 10 mL of Phenol/Chloroform/Isoamyl Alcohol to each tube.
10. Rotate tubes at 40 rpm/min for 10 minutes at room temperature. Then centrifuge at 4,000 xg for 10 minutes.
11. Carefully remove the upper phase by decanting to a fresh 50 mL centrifuge tube, and then add 30 mL of cold 100% ethanol and 4 mL of 5 M ammonium acetate for DNA precipitation. Invert the mixture 20 times to ensure thorough mixing.
12. Centrifuge at 12,000 xg for 5 minutes at 4 °C.
13. Wash DNA pellets with 20 ml 70% ethanol two times and dry the DNA pellet at room temperature for 5 minutes.
14. Resuspend the DNA in 100 μL of 10 mM Tris-HCL (pH 8.0). Transfer resuspended DNA into 1.5 mL DNA LoBind Tubes. Rotate at 10 rpm at room temperature overnight before storing at 4°C for future use.

###### ⚠ CAUTION

High molecular weight (HMW) DNA tends to be very viscous, leading to heterogeneous dissolution during rehydration. To homogenize HMW gDNA, incubate genomic DNA (gDNA) at 50 °C for 1 hour, followed by gentle pipetting using wide-bore tips.

##### Ⓘ TROUBLESHOOTING

15. Use a broad range dsDNA quantitation kit to detect HMW genomic DNA concentration.
16. Assess gDNA purity with OD260/280 by Nanodrop.
17. Check integrity of HMW DNA on a 0.6% 1X TAE agarose gel with 1 kb Extend DNA ladder. For unsheared high-quality HMW DNA, we expect most of the DNA to be larger than 48.5 kb.

###### ⚠ CAUTION

❖ Recommended genomic DNA sample quality before moving forward: unsheared, HMW DNA at ≥210 ng/μL and stored in Tris or TE buffer (pH 8.0) or nuclease-free water.
❖ Whenever possible, use wide-bore tips to minimize shearing of long DNA.

#### crRNA/tracrRNA annealing

18. Preheat the thermal cycler to 95 ºC.
19. For 10x cleavage reactions, sequentially add 0.25 μL each of distinctive crRNA probes (100 uM aliquots in TE buffer) in a 1.5 ml DNA LoBind tube by mixing equal volumes of each crRNA which is dissolved at 100 μM in TE buffer.
20. Mix crRNAs with tracrRNA by adding the following components for every 10 reactions in a 0.2 mL PCR tube:

| Reagent | Volume(μL) for 10x reactions |
| --- | --- |
| 100 μM crRNAs<br>(For FMR1, pool 4 crRNAs) | 1.0<br>(0.25/each) |
| 100 μM tracrRNA | 1.0 |
| Duplex buffer | 8.0 |
| Total | 10 |

21. Pipette to mix the components and then spin them down quickly in a centrifuge.
22. Employ a thermal cycler to incubate the reaction mixture at 95 ºC for 5 minutes. Then, remove mixture from the thermal cycler and cool it down to room temperature over a period of 20 minutes before centrifugation to consolidate any liquid present at the bottom of the tube.

##### ⚠ CAUTION

It is not advisable to store and recycle use of the annealed mixture.

#### Cas9 RNP assembly

23. Thaw NEB rCutSmart Buffer, vortex, and place on ice.
24. To generate Cas9 RNP complexes, combine the elements listed in the table in order, in a 1.5 ml DNA LoBind tube in order for every ten reactions:

| Reagent | Volume(μL) for 10x reactions |
| --- | --- |
| Annealed crRNA with tracrRNA mixture (10 μM) | 10 (Step 22, above) |
| 10x NEB rCutSmart buffer | 10 |
| Nuclease-free water | 78.4 |
| 62 μM SpCas9 Nuclease | 1.6 |
| Total | 100 |

##### ⚠ CRITICAL STEP

Prepare Cas9 cleavage mixture without crRNA•tracrRNA(“Cas9 RNPs minus”) as a negative control to test Cas9 RNP complex cleavage efficiency. Place the components sequentially into a 1.5 ml DNA LoBind tube per 10 reactions:

| Reagent | Volume(μL) |
| --- | --- |
| Duplex Buffer | 10 |
| 10x NEB rCutSmart buffer | 10 |
| Nuclease-free water | 78.4 |
| 62 μM SpCas9 Nuclease | 1.6 |
| Total | 100 |

25. Ensure homogeneous mixing by vigorously flicking the tube.
26. Initiate the formation of Cas9/CRISPR complex by incubating the tube at ambient temperature for 30 minutes, subsequently placing the tubes on ice until further use.

#### HMW genomic DNA dephosphorylation

27. Transfer 5 μg of gDNA into a 0.2 mL PCR tube per reaction. Repeat this step for all samples.

#### ⚠ Important

❖ We recommend including one extra replicate to perform the negative control reaction with Cas9 RNPs minus.
❖ We recommend 2 or 3 replicates for each sample to get more on-target reads.
28. Add nuclease-free water to adjust the volume of each sample to 24 μL, mix thoroughly by gently flicking the tubes. Spin down each sample briefly.
29. Equilibrate the quick calf intestinal phosphatase (CIP) to room temperature prior to utilization. Subsequently, achieve a homogeneous mixture by repetitively pipetting.
30. Add the CIP digestion components in a separate clean 1.5 ml DNA LoBind tube for each 10X reaction:

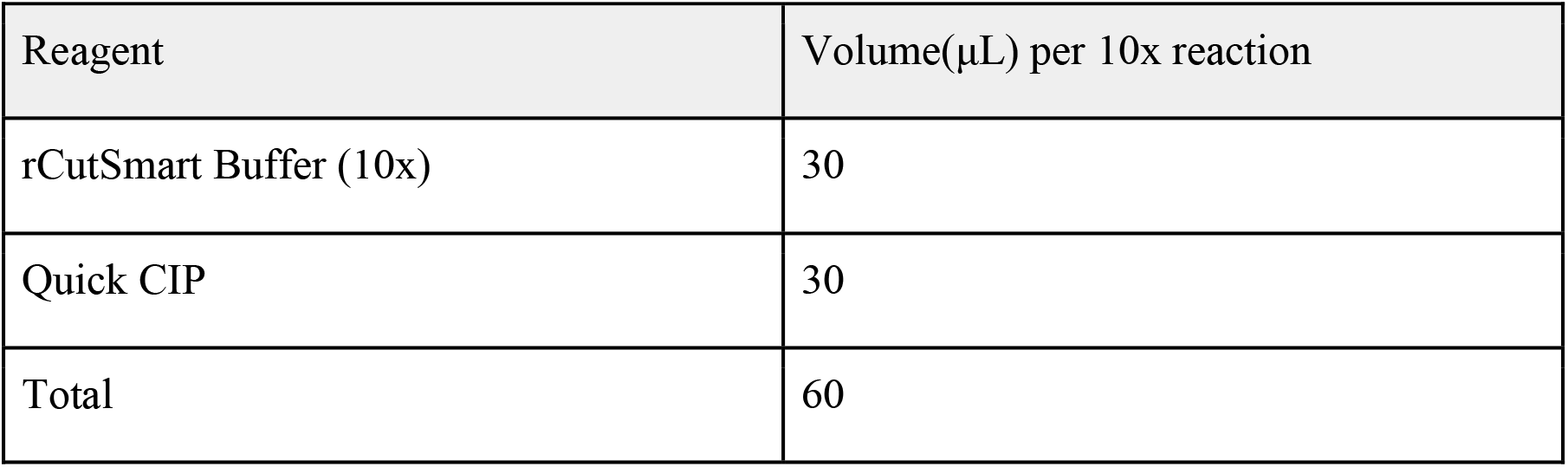

| Reagent | Volume(μL) per 10x reaction |
| --- | --- |
| rCutSmart Buffer (10x) | 30 |
| Quick CIP | 30 |
| Total | 60 |

31. Vortex the mixture, and spin down.
32. Add 6 μL CIP digestion mixture into each prepared sample from step28.
33. Flick the tube gently, and spin down.
34. Employ a thermal cycler to incubate above samples at 37 ºC for 20 minutes, followed by 80 ºC for 2 minutes, then hold at 20 ºC.

#### Cas9 RNP cleavage and dA tailing

35. Thaw 100 mM dATP stock, mix thoroughly by vortex, spin down and put back on ice.
36. Vortex Taq polymerase tube, spin down and put back on ice.
37. To prepare a 10 mM dATP work solution, add 1 μL of the 100 mM dATP stock solution into 9 μL of nuclease free water in a 1.5 ml DNA LoBind tube. Vortex the mixture, spin down and put back on ice.
38. Add the following components into the PCR tube containing 30 μL of dephosphorylated DNA:

| Reagent | Volume(µL) |
| --- | --- |
| Dephosphorylated gDNA (Step 34) | 30 |
| Cas9• RNPs*/Cas9 RNPs minus** | 10 |
| 10 mM dATP | 1 |
| NEB Taq polymerase | 1 |
| Total | 42 |

#### ⚠ CAUTION

*: Use Cas9/CRISPR for HMW gDNA cleavage reactions
**: Use Cas9 without CRISPR as a negative control for each sample
39. Mix the contents within the tube carefully by gentle flick or inversion, followed by centrifugation and subsequent placement in a thermal cycler.
40. Employing the thermal cycler, incubate samples at 37 ºC for 60 mins, followed by 72 ºC for 5 min, and finally, hold at 4 ºC, or place the tube on ice.

#### ⚠ CRITICAL STEP

We recommend initiating Cas9 RNP cleavage at 37 ºC for at least 15 mins, but no more than 60 mins, which may cause more off-target products.
41. Take 1 μL of Cas9 RNPs-cut genomic DNA from each reaction, as well as the negative control, for the PCR template. Set up PCR reactions, as follows:

| Reagent | Volume(µL) |
| --- | --- |
| GoTaq® Green Master Mix (2X) | 12.5 |
| Genomic DNA cut by Cas9 RNPs / Cas9 RNPs minus | 1 |
| Forward primer (10 µM) | 1 |
| Reverse primer (10 µM) | 1 |
| Nuclease-free water | 9.5 |
| Total | 25 |

#### ⚠ CRITICAL STEP

To explore Cas9 RNPs cleavage efficiency, we recommend setting up two distinct PCR reactions: one upstream and the other downstream of the target (see **Box 4**).
42. Run PCR amplification protocol in the thermal cycler with initial incubation at 95 ºC for 5 mins, then 30 cycles: 95 ºC, 2 mins for denaturation; 56 ºC, 30 seconds for annealing; 72 ºC, 90 seconds for extension, followed by a final extension period of 5 minutes at 72 °C, ending with either holding the sample at 4 ºC or returning it to ice.
43. Check Cas9 RNA cleavage efficiency on a 1% 1X TAE agarose gel.

##### Ⓘ TROUBLESHOOTING

###### Purification

44. Transfer Cas9-cut genomic DNA (42 μL) into new 1.5-ml DNA LoBind tube. Add 16 μL of 5 M ammonium acetate and 126 μL of ice-cold ethanol (absolute) and gently invert ten times. Pool replicated samples together.
45. Spin down precipitated DNA at 16,000 xg on centrifuge at 4 °C for 10 min. Wash DNA pellet with 400 μL 70% Ethanol and centrifuge at 16,000 xg for 5 min. Repeat this wash step one more time.
46. Remove the supernatant and dry DNA pellet at RT for ∼5 min.
47. Add 30 μL of 10 mM Tris-HCl (pH 8.0) per reaction, resuspend the DNA pellet gently by flicking the tube or pipetting and incubate them at 50 °C for 1 h. Keep samples at 4 °C for temporary storage, or continue to size selection.

###### BluePippin size selection

48. Program the size selection protocol using the following parameters and save the protocol.
  - Cassette: 0.75DF 3-10 kb Marker S1 Improved Recovery
  - Collection mode: Range (5-12 kb)
  - Reference Lane: 1, “APPLY REFERENCE TO ALL LANES”
49. Calibrate the optics with the BluePippin calibration fixture and check the cassettes before loading samples: Make sure all buffer reservoirs are almost full, avoid using lanes with bubbles or lamination in the agarose, and remove air bubbles behind elution wells.
50. Place the cassette in the optical nest, remove white adhesive strips, and replace the buffer from all elution modules with 40 μL fresh electrophoresis buffer. Seal the elution wells with new adhesive strips.
51. Perform the continuity test by clicking the ‘TEST’ tab.
52. Prepare samples by adding 10 μL loading solution into 30 μL DNA samples and mix samples thoroughly by flicking the tube.

#### ⚠ CAUTION

Recommended sample load guidelines: maximum loading 5 μg/well.
53. Load 40 μL DNA marker S1 into the first well as reference lane 1, and then load each 40 μl DNA sample into the remaining each well.
54. Close the lid, load the protocol name and press “START” to run the program.
55. After running, carefully transfer the contents from each elution well to a 1.5 mL DNA LoBind tube and quantify size-selected DNA samples by Qubit.

#### ⚠ CAUTION

To maximize DNA elution efficiency, we recommend allowing samples to remain in the cassette for 45 mins after the size selection is complete prior to removing eluent.

##### Ⓘ TROUBLESHOOTING

56. Continue to barcoding steps, or keep the size selected samples at 4 °C for <3 days. Alternatively, store samples at -20 °C for long-term storage.

#### ⚠ CAUTION

1. For *FMR1* CGG targeting, the size-selected DNA ranges from 7 kb (normal-length) to 9.5 kb (disease-length).
2. We recommend “Range mode,” which results in higher recovery of desired fragments compared to tight mode and time mode.
3. The degree of target coverage correlates approximately with the quantity of input. When targeting *FMR1* CGG, we generally get 25-50 ng of size-selected DNA from one BluePippin lane with an input of 5 μg HMW gDNA.
4. For achieving maximal target coverage, Cas9 targeted sequencing necessitates a minimum of approximately 5 pg of target material. For example, for the enrichment of an 8 kb human gene, at least 2 μg of high molecular weight genomic DNA would be required as input material.
5. Use “0.75DF 3-10 kb Marker S1 Improved Recovery” instead of the “0.75DF 3-10 kb Marker S1” Cassette Definition. The only difference between the standard and Improved Recovery version is that, in the latter, elution uses direct current rather than pulsed field current leading to higher recovery at the end of the run.

###### Barcoding

57. Thaw the native barcodes at room temperature. Mix each barcode tube by pipetting, spin down and place them on ice. Additionally, warm AMPure XP beads at room temperature for further use.

#### ⚠ CRITICAL STEP

We employ native barcodes from the Native Barcoding Expansion kit, which are ligated to the dA tailed ends. To facilitate multiplexing, a distinct barcode is selected for each sample, enabling their simultaneous run on the same flow cells. In this protocol, we simultaneously barcode 8-12 samples.
58. For each barcode, add reagents in the specified order in a new 1.5 ml DNA LoBind tube:

| Reagent | 1x Reaction (μL) | 3x Reaction (μL) |
| --- | --- | --- |
| Nuclease-free water | 7 | 21 |
| Native Barcode | 3 | 9 |
| Blunt/TA Ligase Master Mix | 50 | 150 |
| Total | 60 | 180 |

#### ⚠ IMPORTANT

To increase output from MASTR-seq and high on-target coverages per sample, it’s recommended to set up 2-3 replicated barcoding reactions according to replicates for each sample.
59. Mix well by vortex or pipetting. Then, spin down the barcode ligation mixtures.
60. Add 60 μL of the native barcode ligation mixture to 40 μL size selected genomic DNA for each reaction (total 100 μL). Mix gently by flicking the tube and spin down.
61. Incubate the barcode ligation reaction for 10 minutes at room temperature.
62. During the incubation time, prepare fresh 5 mL of 70% ethanol in nuclease-free water.
63. Stop barcode ligation by adding 1 μL of 500 mM Tris–EDTA (pH 8.0) (to achieve a final concentration of 5 mM) into each reaction.
64. Start round 1 pooling samples pooling by combining every 3X barcoded reactions (total 300 μL) into a new 1.5 ml DNA LoBind tube. Mix the AMPure XP beads thoroughly by vortex. Immediately add 150 μL of well-resuspended AMPure XP beads to each pooled sample and mix by gentle pipetting.

| Barcoded Reactions | 1~3 | 4~6 | 7~9 | 10~12 |
| --- | --- | --- | --- | --- |
| Number of round 1 pooled tubes | 1x | 2x | 3x | 4x |
| Total volume (μL)/tube* | 300 | 300 | 300 | 300 |
| AMPure XP beads (μL)/tube** | 150 | 150 | 150 | 150 |
| AMPure XP beads (μL) required in total | 150 | 300 | 450 | 600 |

#### ⚠ CAUTION

* When pooling less than 3X barcoded ligation reactions, adjust to total volume of 300 μL with nuclease-free water.
** The volume of AMPure XP beads added is equal to 50 % volume of the pooled reactions per tube. For example, add 150 μL of resuspended AMPure XP beads into 300 μL of pooled 3X barcoded reactions.
  65. Incubate AMPure XP beads with samples for 10 minutes at room temperature without agitation or pipetting.
  66. Spin down the samples and place the tubes on a magnetic rack until the supernatant becomes clear. Carefully remove the supernatant while keeping the tube on the magnet.
  67. Place the tube on the magnetic stand and carefully rinse the beads with 600 μL of freshly prepared 70% ethanol, taking care not to disrupt the pellet. Subsequently, remove the ethanol using a pipette.
  68. Repeat the preceding washing step.
  69. Centrifuge the tube and return it to the magnet. Use a pipette to remove any remaining ethanol. Air-dry the pellet for approximately 30-60 seconds, ensuring not to excessively desiccate it to the point of cracking.
  70. To start round 2 sample pooling, remove tubes from magnetic rack and resuspend pellet by adding recommended volume of nuclease-free water into each tube by following the below table, then combine all the resuspended pellets into one mixture (total of 66 μL). Incubate the mixture for 10 minutes at room temperature.

| Barcoded Reactions | 1~3 | 4~6 | 7~9 | 10~12 |
| --- | --- | --- | --- | --- |
| Number of Round 1 pooled tubes | 1x | 2x | 3x | 4x |
| Nuclease-free water (μL)/tube | 66 | 33 | 22 | 16.5 |
| Number of Round 2 pooled tubes | 1x | 1x | 1x | 1x |
| Total volume after pool (μL)/tube | 66 | 66 | 66 | 66 |

#### ⚠ IMPORTANT

When preparing 2^nd^ samples pooling, we recommend adding volume(=66/N*) of nuclease-free water to 1^st^ step pooled tubes,

*N is number of how many tubes from 1^st^ step pooled.
  71. Use a magnet to pellet the beads until the eluate becomes clear and colorless.
  72. Transfer and preserve approximately 66 μL of the eluate into a new 1.5 ml DNA LoBind tube.
  73. Measure 1 μL of the eluted sample using a Qubit fluorometer. This process leaves approximately 65 μL for adapter ligation.

#### Adaptor ligation

74. Thaw Ligation Buffer and Adapter (AMII) at room temperature, mix each tube thoroughly, by vortexing, and spin down. Subsequently, place them on ice. Meanwhile, warm AMPure XP beads to room temperature. Thaw aliquots of Long Fragment Buffer (LFB) and Elution Buffer (EB) at room temperature.

#### ⚠ CRITICAL STEP

Use AMII adapter from the Native Barcode Expansion 1-12 or 13-24 kits.
75. Add the following components into a new 1.5 ml DNA LoBind Tube:

| Reagent | Volume(µL) |
| --- | --- |
| Ligation Buffer (LNB) | 20 |
| NEBNext Quick T4 DNA Ligase | 10 |
| AMII Adapter | 5 |
| Total | 35 |

#### ⚠ CRITICAL STEP

Ensure the AMII Adapter is added lastly during the ligation step.
76. Mix the ligation mixture thoroughly by pipetting. Since Ligation Buffer (LNB) is highly viscous, ensure a well-mixed adapter ligation mix. Meanwhile, adjust total volume of the multiplexed sample to 65 μL in total by adding nuclease free water.
77. Add 20 μL of adapter ligation mixture into the multiplexed sample (65 μL), and mix gently by flicking the tube and avoiding centrifugation at this stage. Subsequently, promptly add the remaining 15 μL of the adapter ligation mixture to above reaction, resulting in a 100 μL of ligation mixture.
78. Flick the tube gently to mix and then spin down.
79. Incubate the adapter ligation reaction for 10 minutes at room temperature.
80. Vortex to resuspend the AMPure XP beads, making sure they are at room temperature.
81. Add 100 μL 1x TE buffer (pH 8.0) to the adaptor ligation mixture. Gently flick the tube to mix thoroughly.
82. Add 0.4x volume (80 μL) of well resuspended AMPure XP Beads to the ligation mixture. Invert gently to mix. Spin down very gently to keep beads suspended in liquid.

#### ⚠ CRITICAL STEP

When the AMPure beads-to-sample ratio is lower than 0.3:1, DNA fragments of any size will be more easily lost during the clean-up. We calculate volume of AMPure XP beads to be added based on the total volume after addition of TE buffer.
83. Incubate above mixture for 10 minutes at room temperature without agitation or pipetting.
84. Spin down the sample and pellet AMPure XP beads on a magnetic rack until the eluate becomes clear. Maintain the tube on the magnet and pipette off the supernatant.
85. Wash the beads by adding 250 μL of Long Fragment Buffer (LFB). Flick to resuspend beads, return to the magnetic rack, and allow beads to pellet. Subsequently, dispose the supernatant.
86. Repeat the preceding step.
87. Spin down and return the tube to magnetic rack. Remove any residual supernatant by pipette. Air-dry the pellet for 30 seconds, avoiding over-drying to prevent pellet cracking.
88. Remove the tube from magnetic rack, add 13 μL of Elution Buffer (EB) and gently resuspend the pellet by pipetting. Incubate the mixture for 10 minutes at ambient temperature.

#### ⚠ CAUTION

If size of DNA fragments is more than 30 kb, we recommend an incubation time of 30 minutes.
89. Pellet AMPure XP beads on a magnetic rack until the supernatant becomes colorless.
90. Transfer 13 μL of eluate into a new 1.5 ml DNA LoBind tube.
91. Quantify adapted DNA using 1 μL of eluate by a Qubit fluorometer. The remaining 12 μL DNA is used for preparing loading library. Store this sample on ice until ready to use.

#### ⚠ CAUTION

To check long-read DNA library quality, run 2100 bioanalyzer dsDNA high sensitivity DNA assay. We recommend keeping adapter-ligated DNA samples at 4°C for short term storage and storing libraries at -80°C for long term storage.

##### Library loading and sequencing

92. Thaw Flush Tether (FLT), Flush Buffer (FB), Sequencing Buffer (SQB) and Loading Beads (LB) at room temperature, then vortex, spin down and put them on ice after thawing.
93. Open the MinION sequencer Mk 1B lid, slide the MiniON flow cell under the clip, and press down firmly for correct thermal and electrical contact. Connect MinION sequencer to the computer.
94. Perform MinION flow cell QC check by following tutorial from Oxford Nanopore Technologies. Ensure that the number of active pores is above 800.
95. Prepare flow cell priming buffer by adding 30 μL of FLT directly to the tube of FB, and vortex to mix well.
96. Slide the priming port cover clockwise to open the priming port and check for air bubbles under the cover. Remove any bubbles by drawing back a small volume using a P1000 pipette. Verify an uninterrupted buffer flow from priming port to sensor array.
97. Load 800 μL priming buffer into flow cell via the priming port carefully, avoiding any air bubbles. Leave for 5 minutes while preparing the loading library in the following steps.
98. Mix the LB contents thoroughly by pipetting.
99. In a new DNA Lo-Bind tube, prepare the loading library as follows:

| Reagent | Volume($\mu\text{L}$) |
| --- | --- |
| Sequencing Buffer (SQB) | 37.5 |
| Loading Beads (LB) | 25.5 |
| DNA library | 12 |
| Total | 75 |

#### ⚠ CAUTION

Make sure to mix loading Beads (LB) thoroughly and immediately after mixing. As recommended, total volume of library should not exceed 75 μL.
100. To accomplish thorough priming of the flow cell, carefully elevate the SpotON sample port cover to expose the SpotON sample port and add 200 μL of the priming mixture into the flow cell through its designated priming port while ensuring that air bubbles are avoided.
101. Gently mix the loading library by pipetting.
102. Add 75 μL library to the flow cell through the SpotON sample port, loading it drop by drop. Make sure each droplet fully enters the port before introducing another droplet.
103. Carefully place the SpotON sample port cover back, shut priming port, and close the MinION Mk 1B lid.
104. Set up the run settings on the MinKNOW software as follows:

|  |  |
| --- | --- |
| Specified run length | 48 hrs |
| Active channel selection | On |
| Pore scan freq. | 1.5 hrs |
| Reserved pores | On |
| Minimum read length | 1000 bp |
| Basecalling | Off |
| Modified basecalling | Off |

#### ⚠ CAUTION

In this protocol, we always turn off ‘Basecalling’ in our laptop and perform guppy base-calling in high performance computing (HPC).
105. Start the run and sequence for 48 hours.

#### Ⓘ TROUBLESHOOTING

##### Data processing

106. We begin processing raw data (fast5 files) for base-calling using Guppy (version 6.2.1).

guppy_basecaller -i <Test_input_fast5 path> -r -s <basecalled_output_folder path> --config dna_r9.4.1_450bps_hac.cfg --cpu_threads_per_caller <number_of_threads > --num_callers <number_of_callers>

#### ⚠ CAUTION

We have conducted tests using different modes (fast, hac, sup) for base-calling and observed that hac mode is most effective for *FMR1* CGG STR data processing.

##### Ⓘ TROUBLESHOOTING

107. Following base-calling, we demultiplex fastq data for each sample.

guppy_barcoder --input_path <basecalled_output_folder path> --save_path <Debarcoded_output folder path> --config configuration.cfg
108. We align the demultiplexed data to human reference genome hg38 using minimap2 (version 2.22-r1101) to obtain the aligned data (sam files).

minimap2 -a -ax map-ont <Reference folder path>/reference.idx <Debarcoded_output folder path>/sample01.fastq -o <Output folder path>/sample01.sam
109. We developed a tool using python to calculate the number of STRs in each read. Briefly, it takes the FASTQ and minimap2 aligned SAM file as input files. Reads are processed through multiple filtering processes: 1) truncated reads are removed; 2) reads that are not aligned to the targeted regions are removed; 3) reads with anchor regions (i.e. upstream and downstream to CGG tract) and four consecutive CGGs were used for further processing. The number of STRs was extracted by searching the start and end instances of the STR tract. This tool outputs STRs count number for each read in a tab separate file for each sample separately and visualizes the STR counts in a jitter plot and visualizes the reads for all samples using a custom R script with default color schemes. Visualizing the reads is further explained in the step 110.

python MASTRseq.py <FASTQ_files> <SAM_files> <STRAND> <CONFIG_FILE> <DIAGNOSTIC_PLOTS> <OUTPUT_DIRECTORY> Example:

Python MASTRseq.py data/fastq_files/sample_A.fastq,data/fastq_files/sample_B.fastq data/sam_files/sample_A.sam,data/sam_files/sample_B.sam “reverse” config/FMR1_CGG.config “n” out_AB

##### Ⓘ TROUBLESHOOTING

110. We visualized the reads by coloring assigning the individual colors to different bases:

- T – grey
- G – grey
- C – grey
- STR – pink
- N – grey

Rscript scripts/plot_bases.R <INPUT_FILE> <OUTPUT_PNG <COLOR_A> <COLOR_T> <COLOR_C> <COLOR_G> <COLOR_STR> <MAX_READS>
111. CpG DNA methylation in *FMR1* gene promoter
  i. Using the Nanopolish tool, we start by indexing the fast5 files using the ‘Index’ command.
  ii. Subsequently, we use the ‘call-methylation’ command to determine CpG methylation within the specified window ‘chrX:147,902,117-147,960,927’ for FMR1 target loci.
  iii. We determine methylated and unmethylated states based on Log2 likelihood values, considering >0.1 as methylated.
  iv. Then, we calculate the proportion of methylated 19 CpGs in each single-molecule read from each sample.
  v. Finally, we exclude all forward reads from further analysis and use the ‘density’ function in R to visualize the resulting proportions as Kernel Distribution Estimation (KDE) plots.
112. CpG DNA methylation at the STR locus
  i. Using STRique tool, firstly, we index fast5 files with ‘index’ command.
  ii. Then, we then compute DNA methylation status of CGG STRs with ‘count’ command with specific models: ‘r9_4_450bps_mCpG.model’ and ‘r9_4_450bps.model’.
  iii. We filter out all reads with scores <4, which indicate low-quality alignment with flanking regions of the CGG STR tract. We only retain reads with high prefix and suffix scores (> 4) for downstream visualization.
  iv. Subsequently, we calculate the number of methylated CGGs and generate jitter plots using R software.

##### Timing

Steps 1–5, cell culture and cell counting: 1 h

Steps 6–17, HMW genomic DNA preparation and quality assay: 1-2 d

Steps 18–22, crRNA/tracrRNA annealing: 0.5 h

Steps 23–26, Cas9 RNP assembly: 1 h

Steps 27–34, HMW gDNA dephosphorylation: 1 h

Steps 35–40, Cas9 RNP cleavage and dA tailing: 1.5 h

Steps 41–43, Cas9 RNP cleavage efficiency test: 3.5 h

Steps 44–47, Purification: 2 h

Step 48-56, BluePippin size selection: 0.5∼2 d Steps 57–73, Barcoding: 2 h

Steps 74–91, Adaptator ligation: 1 h

Steps 92–105, Library loading and sequencing: 48 h

Steps 106–108, Data processing: 12-24 h

Steps 109–110, STR processing and visualization: 2 h

Steps 111–112, DNA methylation analysis: 12-24 h

BOX 2. Quality control assays for high molecular weight genomic DNA: 3 h

BOX 4. Cas9 RNP cleavage efficiency assay: 3 h

BOX 5. Bioanalyzer profile of a final MASTR-seq library: 1 h

##### Troubleshooting

Troubleshooting advice can be found below:

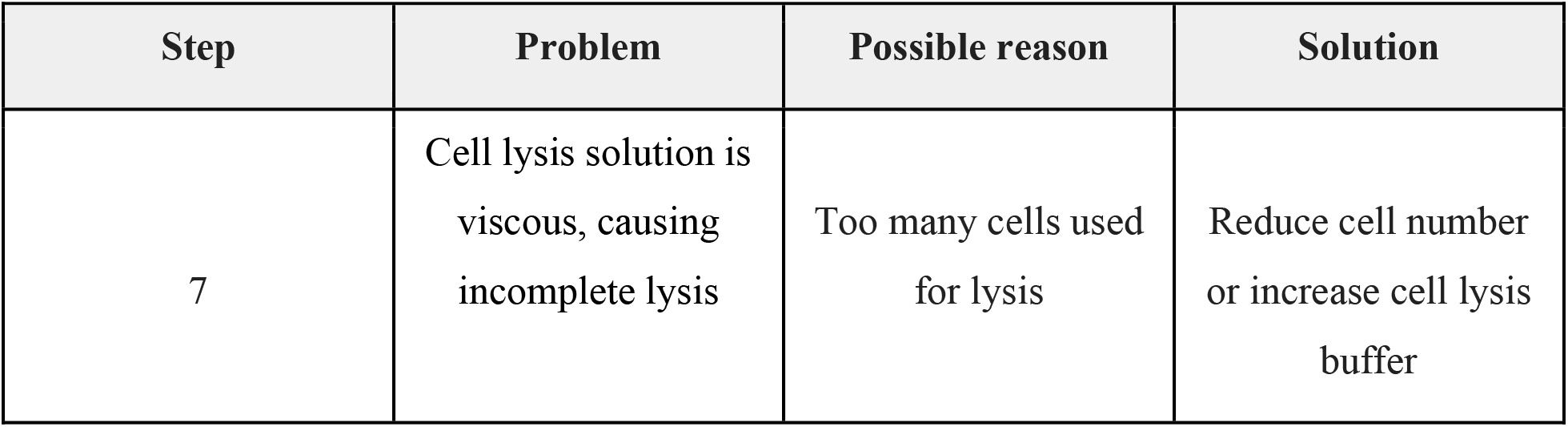
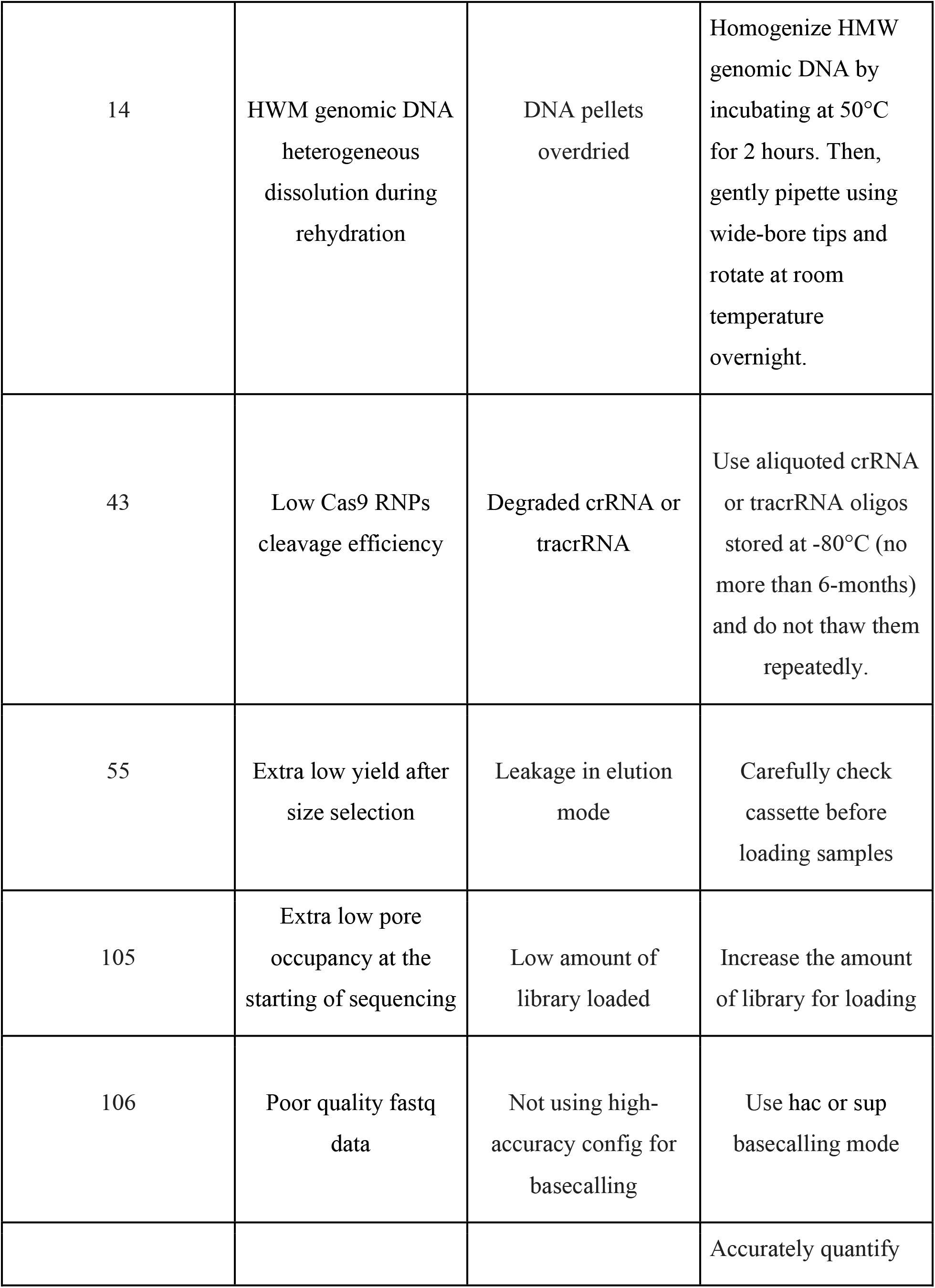
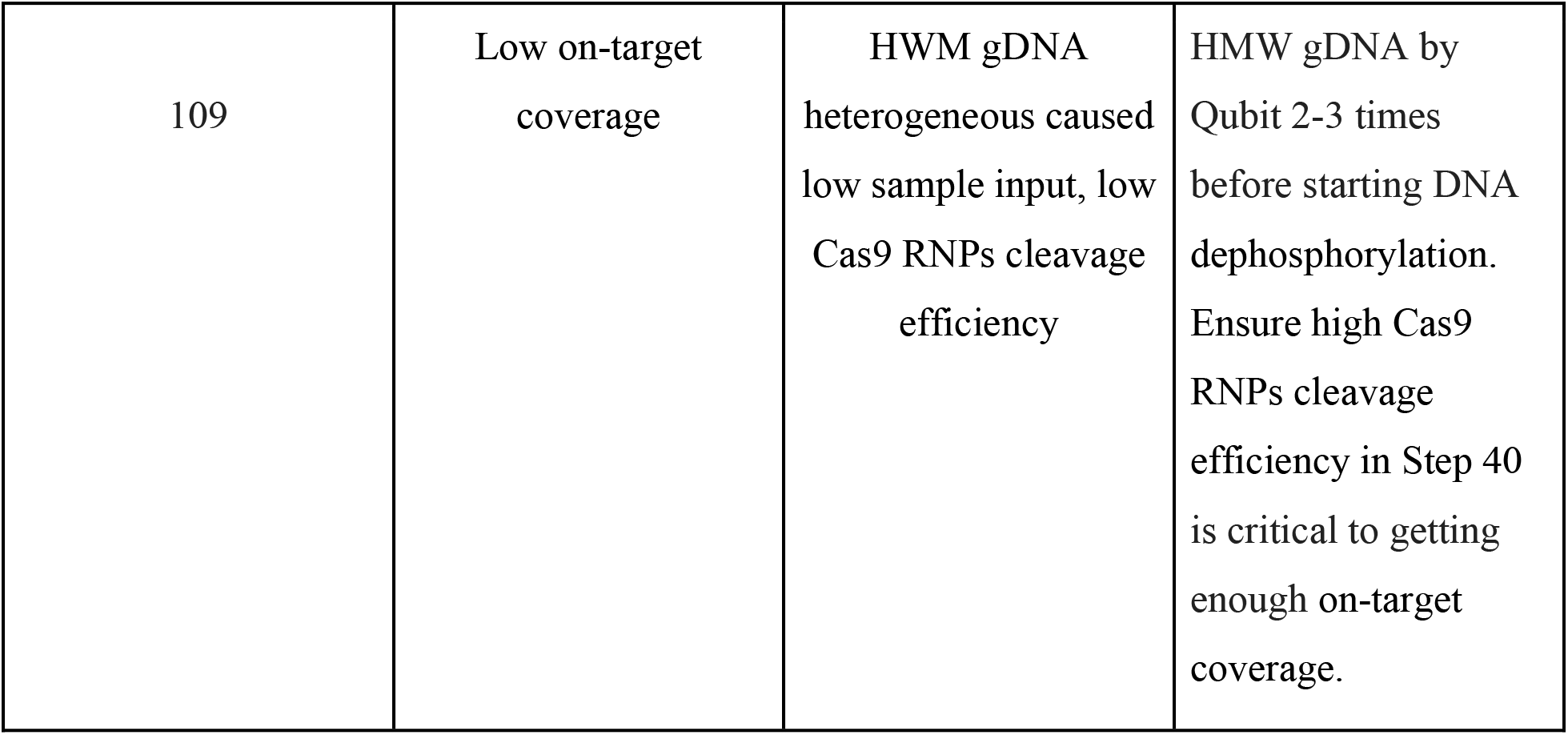

### Anticipated results

To assess the suitability of MASTR-seq for accurate STR length quantification, we quantified the *FMR1* CGG STR across a spectrum of tract lengths including normal-length, premutation-length, and full mutation-length hiPSCs (**Figure 2a**). Implementation of MASTR-seq for STR genotyping yielded over 300x on-target coverage and a >10-fold increase in on-target read proportion (**BOX 6**). We find that forward strand reads have high error rates compared to reverse reads (**Figure 2b-e, Table 8**). We verified that reverse reads represent accurate measurements of *FMR1* CGG STR length by visualizing the reads (**Figure 2b+d, Table 8**). Normal- and premutation-length iPSC lines had, on average, approximately 19-25 and 137 CGG triplets, respectively, whereas all three independent iPSC lines derived from FXS patients showed a similar median of 420-470 CGG triplets (**Figure 2c)**. Furthermore, we employed MASTR-seq to determine the length of the *FMR1* CGG STR tract in premutation-length clones CRISPR engineered by cutting back the mutational-length CGG STR from a parent FXS iPSC line (**Figure 2f-g**). Thus, MASTR-seq efficiently measures the CGG tract length of normal-length, premutation-length, mutation-length, and CRISPR engineered premutation-length cutback iPSC clones.

We demonstrate that nanopore long-read sequencing can accurately quantify STR length and DNA methylation at single molecule resolution. Our method accurately confirms the known hypomethylation of CpG dinucleotides in the *FMR1* promoter in normal-length and premutation-length iPSCs, as well as promoter hypermethylation in mutation-length iPSCs (**Figure 3a**). Moreover, at the CGG STR tract, we observed nearly all hypomethylated alleles in all 3 normal-length and the premutation-length iPSC lines as well as a high level of hypermethylation in mutation-length iPSCs (**Figure 3b)**. Consistent with the DNA methylation data, our published *FMR1* gene expression data revealed high expression in normal-length and premutation-length iPSCs and silencing in mutation-length iPSCs (**Figure 3c**). In CRISPR engineered clones, we demonstrated that 5 out of 7 premutation-length cutback clones exhibit depleted CpG methylation at the *FMR1* promoter and the CGG STR tract (**Figure 3d-e, Table 9-10**). *FMR1* de-repression occurs in the same clones where DNA methylation is depleted compared to the parent FXS patient-derived mutation-length iPSC line (**Figure 3f**). Together, our data demonstrate robust quantification of both DNA methylation and STR length in the same single molecule across a cohort of normal-length, premutation-length, and mutation-length FXS iPSC lines.

To validate MASTR-seq’s utility beyond the CGG STR, we also employed our method to verify the length of a *HTT* CAG tract in three isogenic human embryonic stem cells (hESCs) engineered to model Huntington’s disease (HD) ^63^ (**Figure 4a**). The REUS2 hESC lines have been heterozygously engineered with specific, fully validated CAG tract lengths as indicated by the labels (**Figure 4a**). MASTR-seq accurately confirmed the presence of the unedited allele as well as the NL or ML edited CAG STR tract in the *HTT* gene in all isogenic hESCs (**Figure 4b-c**).

Conventional gold-standard STR genotyping methods employed in clinical settings, such as triplet repeat primed PCR (TP-PCR), suffer from low sensitivity and accuracy in measuring mutation-length STR expansion at single-molecule resolution. We employed the AmplideX mPCR *FMR1* kit to quantify CGG STR length across iPSC lines with NL, PM, FXS and PM cut-back alleles using clinical gold-standard methods (**Figure 5**). As previously reported, TP-PCR is useful only to roughly inform an assessment of premutation- or full mutation-length, but the individual single allele information and accurate tract length genotyping cannot be achieved (**Figure 5a**). By contrast, MASTR-seq demonstrated sensitivity and single-molecule resolution in multimodal measurement of STR length and DNA methylation (**Figure 5b**). Altogether, we demonstrate that MASTR-seq allows for high-throughput, efficient, accurate, and cost-effective measurement of STR length and DNA methylation in the same single allele for up to 8-12 samples in parallel in one Nanopore MinION flow cell. Using Cas9-mediated target enrichment and PCR-free, multiplexed nanopore sequencing, MASTR-seq achieves a >10-fold increase in on-target read proportion for highly repetitive, technically inaccessible regions of the genome relevant for human health and disease.

**Figure 5.**
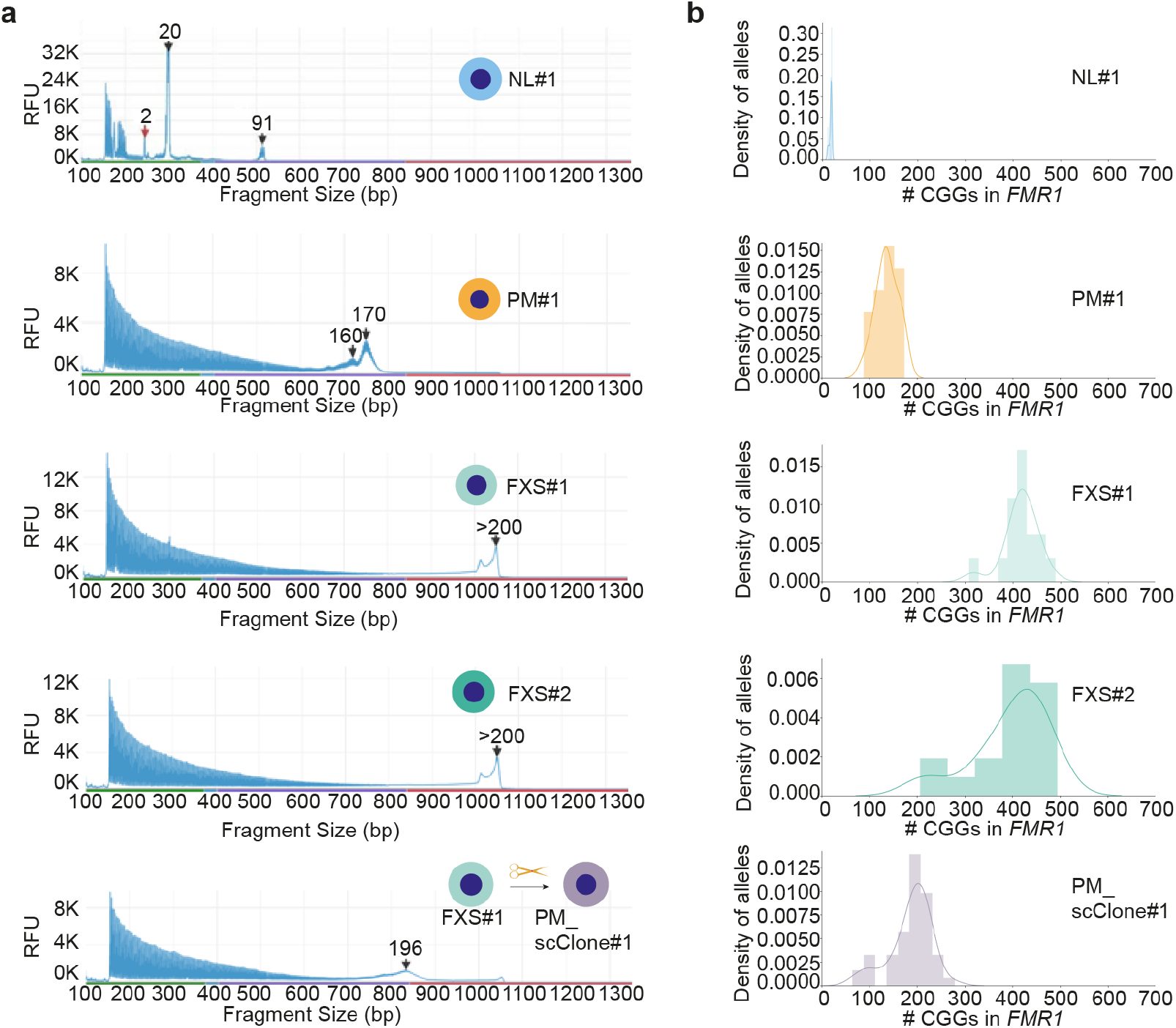
Comparation of PCR based assay and MASTR-seq. **(a)** Capillary gel electrophoresis traces using AmplideX mPCR *FMR1* kit for quantification of *FMR1* CGG STR length among iPSC lines (see Figure 2) with NL, PM, and FXS alleles and PM cut-back single cell clones. **(b)** Histogram and Kernel Distribution Estimation (KDE) of STR tract length from the *FMR1* locus across the same iPSC lines using MASTR-seq.

## Author contribution statement

Conceptualization, C.S., and J.E.P.-C.

Experimentation. C.S., T.M., R.B.

Computation & Visualization, C.S., K.R.C.

Funding, J.E.P.-C.; T.M.

Administration, J.E.P.-C.

Writing, C.S., and J.E.P.-C.

Review & Editing, C.S., K.R.C., T.M., R.B., H.-S.R., and J.E.P.-C.

## Acknowledgments

We thank the members of the Cremins lab for helpful feedback.

## Data availability

All MASTR-seq raw data and processed data have been provided in **Table 11**. To review GEO accession GSE265944, enter private reviewer token: qfmvmmkqfpuprmf.

## Code availability

Code for the figures in this study are available from freely available code in bitbucket: https://bitbucket.org/creminslab/mastrseq

### Related links

**Key reference using this protocol**

Malachowski,T. et al. *Cell* **186**(26),5840-5858.e36(2023). Doi: 10.1016/j.cell.2023.11.019.

