## Supplemental Methods for "MASTR-seq: Multiplexed Analysis of Short Tandem Repeats with sequencing"

Supplementary Information

Chuanbin Su[1,2,3], Keerthivasan Raanin Chandradoss[1,2,3], Thomas Malachowski[1,2,3], Ravi Boya[1,2,3], Han-Seul Ryu[1,2,3], Kristen J. Brennand[4,5,6,7], Jennifer E. Phillips-Cremins[1,2,3,#]

1. Department of Bioengineering, University of Pennsylvania, Philadelphia, PA
2. Epigenetics Institute, Perelman School of Medicine, University of Pennsylvania, Philadelphia, PA
3. Department of Genetics, Perelman School of Medicine, University of Pennsylvania, Philadelphia, PA
4. Nash Family Department of Neuroscience, Icahn School of Medicine at Mount Sinai, New York, NY 10029
5. Department of Genetics and Genomics, Icahn School of Medicine at Mount Sinai, New York, NY 10029
6. Friedman Brain Institute, Black Family Stem Cell Institute, Pamela Sklar Division of Psychiatric Genomics, Icahn School of Medicine at Mount Sinai, New York, NY 10029
7. Department of Psychiatry, Yale School of Medicine, New Haven, CT, 06520

**Supplementary Tables**

Supplementary Table 1. CRISPR RNA design and sequence

Supplementary Table 2. Size selection program.

Supplementary Table 3. Nanopore sequencing setup

Supplementary Table 4. Sequencing summary file

Supplementary Table 5. Barcoding summary file

Supplementary Table 6. On target coverage analysis

Supplementary Table 7. On-target assay among different methods

Supplementary Table 8. CGG count calculated by MASTR_seq.py

Supplementary Table 9. *FMR1* promoter DNA methylation detected by Nanopolish

Supplementary Table 10. *FMR1* CGG methylation quantified by STRique

Supplementary Table 11. Data source link

#### Supplementary Methods

##### On-target coverage calculation

We use samtools to calculate average coverage of *FMR1* target region. The coverage or depth is the average number of times a nucleotide is represented by a high-quality base[1]. Average coverage of target region is defined as the average of coverage (reads number) of all bases between the innermost gRNA sites[2]. On target reads are defined as any read that contains at least 1bp overlap with a target region. Off target reads contain no target-overlapping bases.

##### Public data processing

We downloaded the raw data of Cas9 target nanopore sequencing from the BioProject ID PRJNA531320[52].The fast5 files were retrieved from the run SRR1104786 provided access link. We processed raw data (fast5 files) for base-calling using Guppy (version 6.2.1) with default parameters. We aligned the FASTQ files on the reference genome hg38 using minimap2(version 2.22-r1101) with default parameters and saved the alignments in SAM file format. We used 'Samtools' functions 'view', 'sort', and 'index', respectively, to convert SAM into BAM file format, sorted the reads based on their coordinates and indexed BAM files. We obtained the mean coverage of target regions and on-target reads number using the 'Samtools' functions 'coverage' and 'depth' with the default parameters, respectively. We calculated 'total-aligned' reads using 'Samtools' command 'view'.

Finally, we calculated on-target reads percentage by following formula:
on-target reads percentage = (on-target reads number / total-aligned reads number)*100%

We calculated off-target reads percentage by following formula:
off-target reads percentage = (1-(on-target reads number / total-aligned reads number))*100% .

##### CGG PCR assay

We use AmplideX PCR/CE FMR1 Reagents to quantify *FMR1* CGG STR in human iPSCs.
